# Reconstructing tissue culture to improve Agrobacterium-mediated transformation of maize

**DOI:** 10.64898/2026.05.19.726338

**Authors:** Seijiro Ono, Misato Ono, Reinhold Brettschneider, Dagmar Sauer, Katja Müller, Martina Balboni, Max van der Heide, Arp Schnittger

## Abstract

The ability to insert, delete, or modify genetic information is crucial for mechanistic studies and biotechnological applications. However, efficient genetic transformation remains a major bottleneck for research in maize and many other crops. Here, we report an optimized *Agrobacterium*-mediated transformation platform based on systematic reconstruction of tissue culture handling in the maize inbred line A188. Refinement of callus induction, selection, and regeneration substantially improved recovery of transgenic plantlets. To distinguish independent T-DNA insertion events, we developed TAFLP (T-DNA Amplified Fragment Length Polymorphism), a simple and inexpensive assay that amplifies T-DNA flanking sequences and can be performed using standard laboratory equipment. Our enhanced transformation pipeline was also applicable to the inbred line B104 as well as to hygromycin and G418 selection systems, demonstrating broad utility of our method. We validated the platform for CRISPR/Cas9 mutagenesis and reporter line generation. Using this approach, we isolated new loss-of-function alleles of *MAC1* and *ACOZ1* and generated reporter lines for analysis of meiotic protein dynamics. Together, these results provide a broadly applicable framework for improving maize transformation efficiency and recovering independent transgenic and genome-edited events.

## Introduction

Agrobacterium-mediated transformation remains one of the most important tools in plant biotechnology, enabling stable introduction of transgenes and genome-editing constructs into crop species (Hwang et al., 2017). In maize, however, transformation efficiency is still strongly influenced by several factors, most notably genotype, embryo developmental stage, and tissue-culture conditions (Kausch et al., 2021). Although substantial progress has been made, the recovery of independent transformants is often limited by low embryogenic callus formation, genotype-dependent regeneration capacity, and loss of transformed events during prolonged selection and regeneration. These constraints are particularly problematic when the objective is to generate large numbers of independent events for functional genomics, transgene expression studies, or CRISPR/Cas9-based mutagenesis.

Earlier transformation studies using *Agrobacterium*-mediated transformation and biolistic micro-particle bombardment achieved higher efficiencies with Hi-II hybrid embryos (Armstrong et al., 1991; Zhao et al., 1998; Miller et al., 2002; Vega et al., 2008). This has been attributed to the high capacity of Hi-II embryos to induce type II somatic embryogenic callus, a friable, fast-growing, and highly regenerative callus type (Vega et al., 2008). However, transformation using Hi-II embryos is not ideal for functional genomic studies, primarily because of trait segregation in the offspring. Many phenotypic, physiological, and quantitative trait studies require a nearly uniform genetic background between the established line and control plants, as well as controlled growth conditions. Therefore, transgenic plants derived from Hi-II hybrids generally need to be backcrossed before phenotypic assays can be performed, a process that often takes several years.

A major breakthrough in overcoming genotype dependency was the introduction of the maize *BABY BOOM*(*BBM*) and *WUSCHEL2*(*WUS2*) morphogenic regulators (MRs) (Lowe et al., 2016). In this system, ectopic and/or constitutive expression of MR genes enhances embryogenic callus formation and regeneration in transformed cells. Today, multiple transformation protocols incorporating MRs are available across a broad range of genotypes, and these approaches have led to substantial improvements in the recovery of primary regenerants (Lowe et al., 2018; Mookkan et al., 2018; Hernandes-Lopes et al., 2023; Aesaert et al., 2022). However, overexpression or ectopic expression of MRs often causes abnormal plant development and reduced fertility (Lowe et al., 2016). One widely used strategy to avoid these undesirable developmental effects is Cre/LoxP-mediated excision, in which Cre recombinase is expressed under an inducible promoter, such as the drought-inducible promoter *rab17* (Lowe et al., 2016), the heat-shock-inducible promoters *Hsp17.7* and *Hsp26* (Wang et al., 2020), or the ABA-inducible promoter *pZmGLB1* (Aesaert et al., 2022). Cre then excises the MR and Cre cassette flanked by two LoxP sites. However, this system introduces an additional level of uncertainty during tissue culture, because successful MR removal depends on tissue exposure to the appropriate stress condition to induce Cre expression, which may affect plant recovery and survival. Furthermore, MR and Cre expression cassettes occupy a substantial portion of the T-DNA, while transgene delivery efficiency is influenced by insert size (Xi et al., 2018). This is a considerable limitation, especially when introducing maize reporter constructs, which are often larger because of extended cis-regulatory elements such as promoters, introns, and terminators (Marand et al., 2025; Staut et al., 2025).

Maize inbred lines such as A188 and B104 have been widely used for Agrobacterium-mediated transformation because of their relatively high embryogenic potential (Ishida et al., 2007; Raji et al., 2018). For functional genomic studies, B104 is advantageous because it is phylogenetically closer to B73, the standard maize inbred line widely used in genetic studies and supported by extensive genome annotation. A188, on the other hand, offers advantages for genetic studies because of its relatively shorter life cycle compared with B73 or B104, as well as the availability of a high-quality genome assembly and annotation (Lin et al., 2021). Nevertheless, even in these genotypes, transformation outcomes remain variable and are highly dependent on technical aspects of tissue-culture handling. Conventional protocols often rely on long selection periods, strong antibiotic or herbicide pressure, and limited control over callus morphology and regeneration dynamics. As a result, the number of regenerated plantlets does not always reflect the number of independent transformation events, and chimerism or redundancy can complicate downstream analysis. These issues are particularly relevant for CRISPR/Cas9 experiments, where independent mutant events are often required to recover rare allelic combinations or to establish null mutant lines for essential genes.

Here, we present an optimized maize transformation platform based on the systematic refinement of tissue-culture handling. Using A188 as the primary system, we identified eight key modifications that improved callus induction, selection, and regeneration. To better evaluate the number of distinct transformation events recovered by the system, we also developed TAFLP (T-DNA Amplified Fragment Length Polymorphism), a simple and inexpensive assay for event discrimination based on amplification of T-DNA flanking regions. We demonstrate the utility of this platform in CRISPR/Cas9 mutagenesis and reporter-line generation through two case studies: one targeting the meiosis-related gene *MAC1* and the other targeting *ACOZ1*, a gene required for meiotic chromosome organization.

## Results

### Reconstructing tissue culture methods: eight key modifications for improved maize transformation

To improve maize transformation efficiency through optimized tissue-culture handling, we carried out a series of optimization experiments using the A188 line, an elite maize inbred line with a fully annotated genome and, importantly, an autonomous and high capacity for embryogenic callus induction from immature embryos (Ge et al., 2022). We used the BASTA® (glufosinate) selection system, as in previous studies (Ishida et al., 2007). A detailed step-by-step procedure is provided in the Supplementary Procedure. Here, we describe eight key modifications relative to the conventional method.

First, we used meropenem trihydrate (meropenem) as an alternative to carbenicillin, cefotaxime or vancomycin, to suppress Agrobacterium growth. Meropenem was continuously supplied in all media after inoculation and co-culture (Fig. 1A). Meropenem has previously been reported to be effective at low concentrations without phytotoxic effects (Sjahril and Mii, 2006). Under our working conditions, maize embryogenic callus induction and growth were improved with 25 mg/L meropenem compared with the conventional medium containing 100 mg/L carbenicillin (Supplementary Fig. 1).

**Figure 1.**
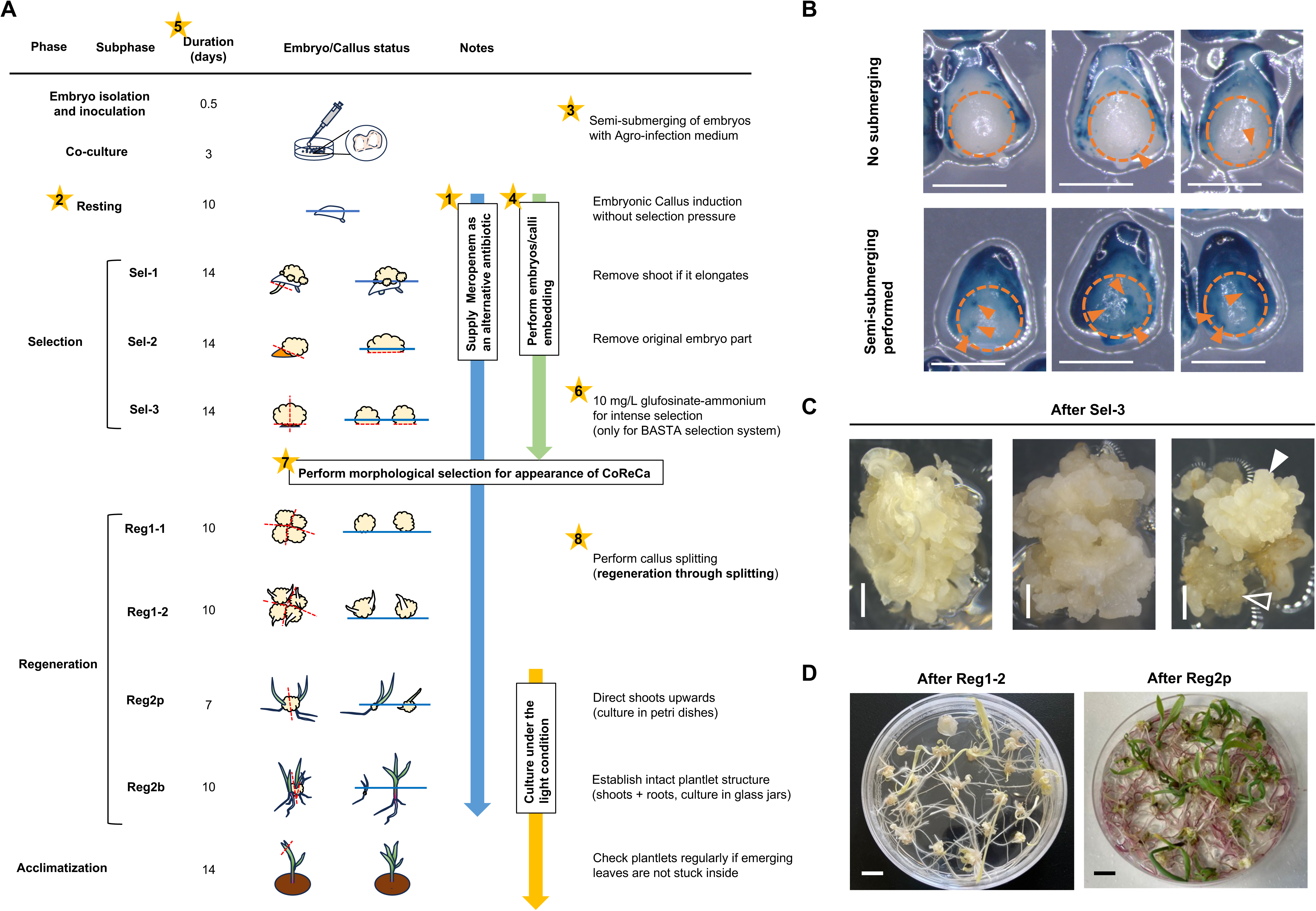
Reconstructed tissue culture handling for Agrobacterium-mediated transformation of the A188 inbred line. A. Overview of the reconstructed tissue culture workflow. Stars with numbers correspond to the major key modifications described in the main text. Blue lines indicate the medium surface relative to embryo/callus embedding, while red dashed lines indicate positions at which removal or callus splitting was performed in the respective phases. B. Comparison of infection events without (upper) or with (lower) embryo semi-submerging during the co-culture phase, monitored by GUS staining of the *UBIpro:GUS* construct. Dashed circles indicate the expected region of embryonic callus induction. Arrowheads indicate positive GUS spots overlapping with the callus-inducing region. Bars = 1 mm. C. Examples of Committed Resistance Callus (CoReCa) morphology after the Sel-3 phase. Closed and open arrowheads indicate CoReCa-positive and CoReCa-negative sectors, respectively, within a chimeric callus. Bars = 1 mm. D. CoReCa-derived calli showing improved regeneration ability after callus splitting. Bars = 1 cm.

Second, we introduced a 10-day resting phase, during which inoculated embryos were cultured on callus-inducing medium without selection pressure, to facilitate embryogenic callus formation (Fig. 1A). A resting phase has been used in particle bombardment-based transformation procedures but is rarely applied in Agrobacterium-mediated methods (Brettschneider et al., 1997; Raji et al., 2018). We hypothesized that this phase increases the amount of callus available for subsequent selection and thereby improves the recovery of transformed cells.

Third, we semi-submerged embryos in liquid Agrobacterium infection medium during the co-culture phase (Fig. 1A). This modification was based on our observation that, without submergence, GUS-positive spots from the *UBIpro:GUS* construct were detected mainly at the edge of the immature embryo scutellum and only occasionally in the upper region of the scutellum (Fig. 1B). In contrast, embryogenic callus arose from the upper scutellar region rather than from the edge (Fig. 1B, dashed circles). For successful recovery of transformants, Agrobacterium infection and callus induction must occur in the same tissue region. Semi-submergence slightly increased infection events in the upper scutellum (Fig. 1B, arrowheads).

Fourth, we defined the degree of embedding of inoculated embryos/calli in each culture phase, as this likely affects nutrient uptake and selection efficiency (Fig. 1A). In particular, slight embedding of embryos during the resting phase was critical for improving embryogenic callus formation (Supplementary Fig. 3). We presume that embedding the embryo suspensor in the medium enhances nutrient uptake, which is required for callus induction while suppressing unwanted shoot primordium development that might otherwise lift the embryo away from the medium. In the later selection phases, the calli were half-embedded to allow direct uptake of nutrients and selective agents from the callus surface (Fig. 1A).

Fifth, we shortened the duration of each tissue-culture phase compared with the conventional method, limiting each phase to a maximum of 2 weeks (Fig. 1A). This was particularly important for the BASTA/glufosinate selection system. In the presence of glufosinate, non-resistant cells metabolize the compound and release ammonium, whereas phosphinothricin acetyltransferase, encoded by the bar gene, detoxifies glufosinate by acetylation (Supplementary Fig. 4A) (Rojano-Delgado et al., 2014). Accumulation of non-resistant cells may therefore adversely affect the growth of resistant cells by altering the culture pH (Supplementary Fig. 4B).

Sixth, we revised the concentration and formulation of the selective agent and used 10 mg/L glufosinate-ammonium salt instead of BASTA® solution for the third selection phase (Sel-3). This enabled strong selection while avoiding the excess surfactant present in BASTA® formulations, which may negatively affect overall callus growth.

Seventh, we established clear morphological criteria for callus selection, especially after the intense Sel-3 phase. We defined post-Sel-3 calli capable of regenerating numerous plantlets in subsequent regeneration phases as committed resistant callus (CoReCa). CoReCa was characterized by: (1) a semi-transparent or pale-white to pale-yellow color; (2) a semi-hard, slightly watery texture, but not extremely hard or dry; (3) extensive folding and budding, sometimes with small hair-like structures resembling leaf primordia on the surface; (4) a solid interior rather than a hollow structure, but one that could be easily split with a razor blade; and (5) a volume at least threefold greater than at the start of Sel-3, suggesting logarithmic-phase cell growth. We considered the appearance of CoReCa to be a major indicator of transformation success after the Sel-3 phase (Fig. 1A,C).

Eighth, we implemented frequent callus splitting, mainly after Sel-3 and during the regeneration phases, and in some cases during the transition from Sel-2 to Sel-3 (Fig. 1A). We refer to this concept as regeneration through splitting. We observed that CoReCa did not regenerate shoot primordia efficiently when maintained as large callus chunks; instead, the calli tended to continue expanding. In contrast, callus splitting generated numerous shoot and root primordia, presumably by increasing callus surface area and improving phytohormone exchange, thereby stimulating regeneration (Fig. 1D). For splitting, calli larger than 10 mm in diameter were cut into pieces smaller than 5 mm in diameter using a razor blade. Splitting should be performed at each regeneration interval using a clean cutting edge rather than by tearing with forceps.

### Boosted transformation efficiency with the revised method

Using the improved tissue culture handling described above, we tested transformation efficiency in 10 independent experiments. For transgene delivery, we developed several reporter constructs derived from maize genomic DNA fragments as well as several CRISPR/Cas9 constructs targeting endogenous maize genes (see Supplementary table 1 and Supplementary procedure for details). We obtained transformation efficiencies ranging from 17.9% to 171.4%, calculated as the number of plantlets regenerated per number of embryos inoculated (Table 1).

**Table 1.**
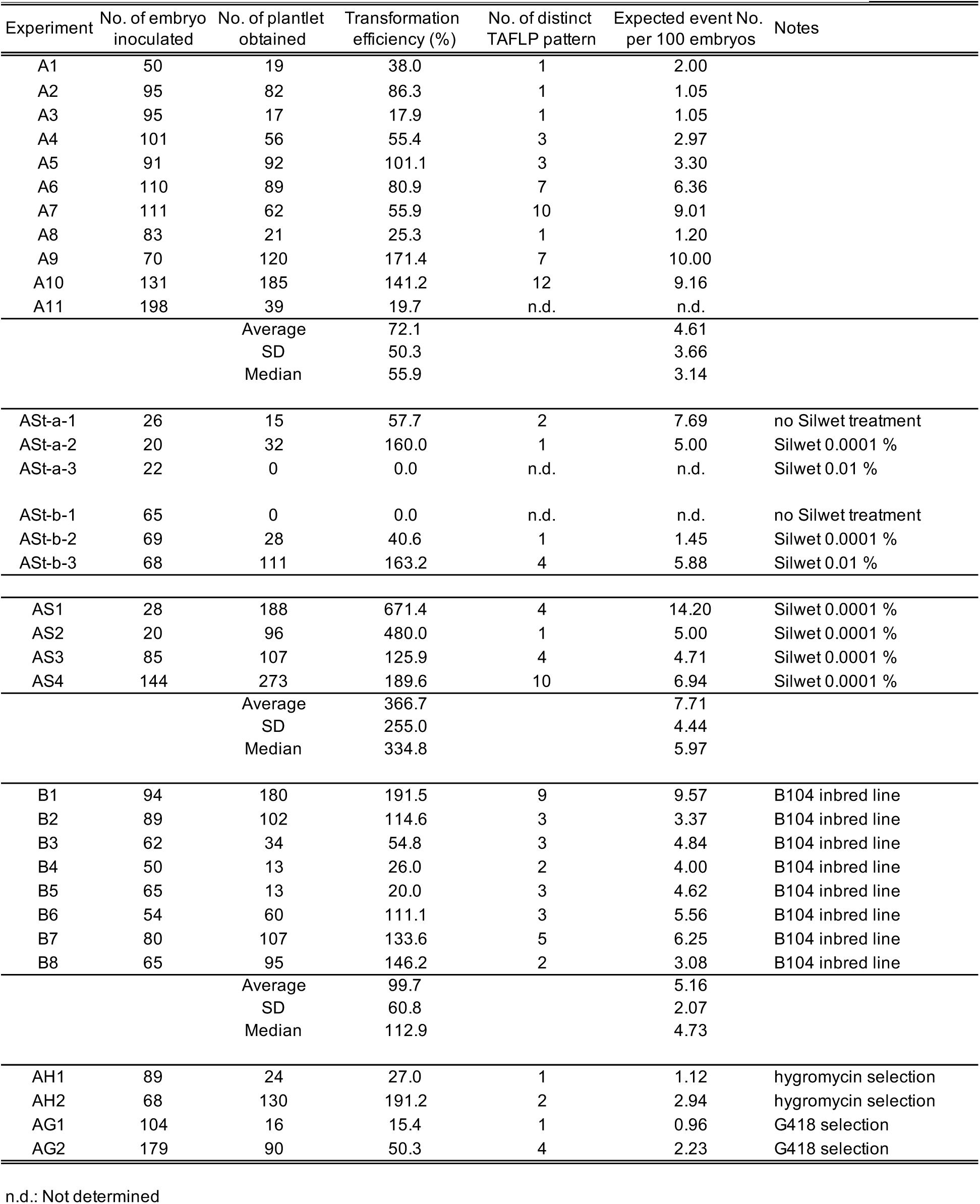
Statistical summary of transformation experiments performed in this study.

Regeneration through splitting was critical for obtaining regenerants efficiently; however, this approach was expected to yield multiple plantlets derived from the same T-DNA insertion event. Tracing the origin of all calli was highly labor-intensive, and random selection of calli during the selection and regeneration phases was not desirable because it could reduce recovery of rare events. In addition, it was not realistic to grow and propagate all regenerated plantlets in the limited greenhouse space available. Therefore, we sought to determine how many independent T-DNA insertion events could be obtained with this method.

To this end, we traced the number of calli and/or plantlets derived from single CoReCa to determine how many plantlets could be regenerated from a single CoReCa appearing at the Sel-3 phase. In experiment A2, two CoReCa appeared at the end of the Sel-3 phase and were cultured as distinct subpopulations during the regeneration phases, resulting in 27 and 43 plantlets, respectively (Supplementary Fig. 5). Another 12 plantlets had been obtained from other calli with incomplete CoReCa morphology in this experiment. Later, we found that all 82 plantlets obtained in this experiment shared the same T-DNA insertion event (see below), indicating that the two CoReCa and the other positive calli originated from a single transformation event and had already undergone callus splitting before the Sel-3 phase (Fig. 1A, Supplementary Fig. 5). Although a single CoReCa may not always represent a single T-DNA insertion event, the number of CoReCa can serve as a useful parameter for estimating the number of independent events in the population.

### TAFLP: an easy, reproducible, and inexpensive assay to distinguish independent T-DNA integration events

Next, we developed an assay to distinguish independent T-DNA insertion events among regenerated populations. Specifically, we sought an assay that could: (1) be performed and analyzed during the vegetative stage of regenerants, i.e. within 2 weeks after acclimatization; (2) require only standard laboratory equipment, such as thermal cyclers and agarose gel electrophoresis systems; (3) be suitable for large populations; and (4) be highly reproducible, yielding the same result among plantlets derived from the same transgenic event. We therefore established TAFLP (T-DNA Amplified Fragment Length Polymorphism), based on the AFLP principle (Vos et al., 1995), which fulfilled all of these requirements (Fig. 2A; Supplementary Procedures).

**Figure 2.**
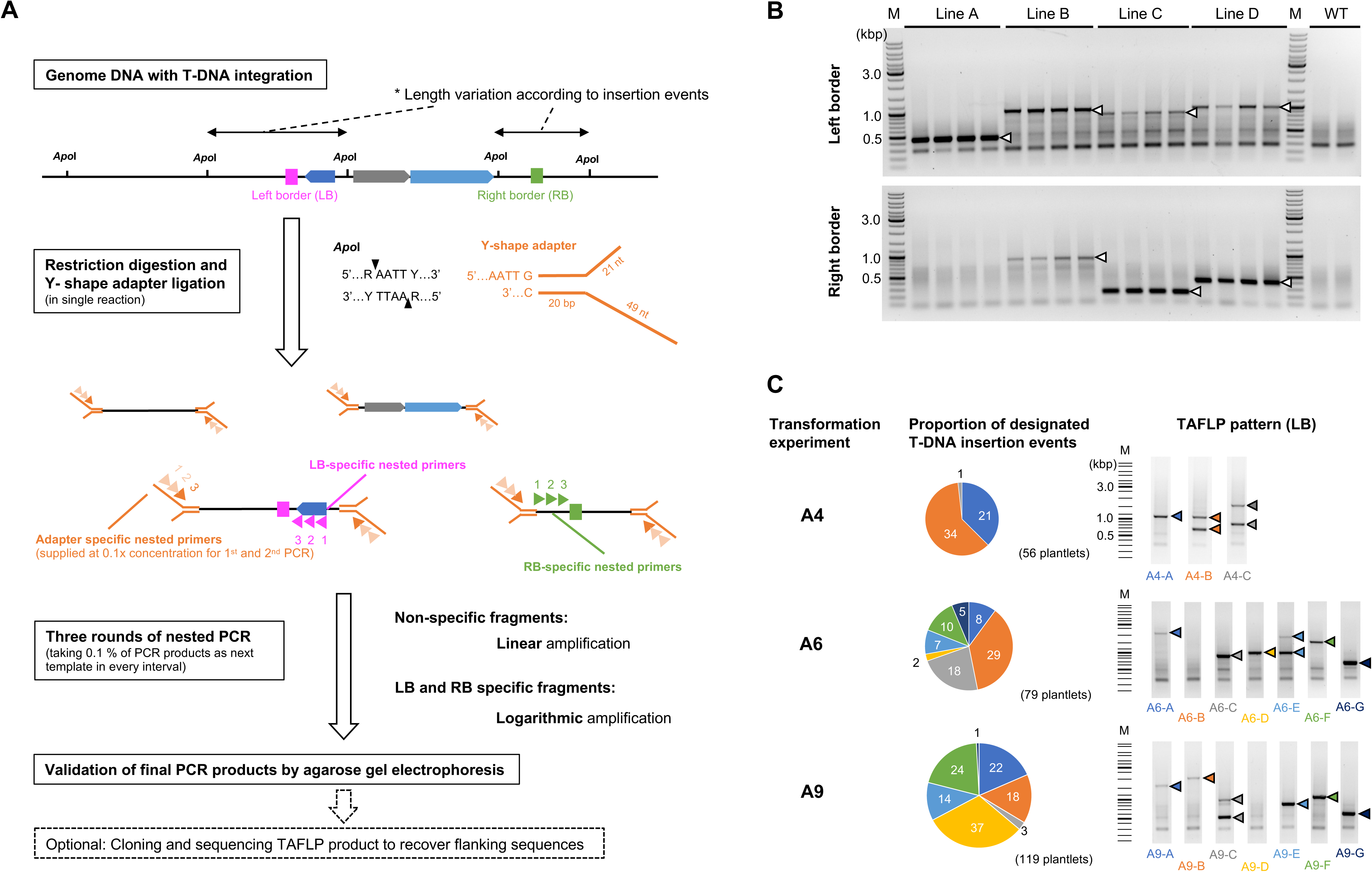
TAFLP assay to distinguish independent T-DNA insertion events. A. Schematic overview of the principle and workflow of the T-DNA Amplified Fragment Length Polymorphism (TAFLP) assay. B. Test experiment using T1 plants from four previously established independent lines (Lines A–D) and wild-type (WT) controls. Arrowheads indicate specific amplification patterns in the respective lines. M, DNA ladder marker (GeneRuler DNA Ladder Mix, Thermo Scientific). C. Proportions of designated T-DNA insertion events in transformation experiments A4, A6, and A9. Numbers in the pie charts represent the number of plantlets in each event. Arrowheads indicate specific amplification patterns for the respective events, with colors corresponding to the pie charts. M, virtual DNA ladder marker.

Briefly, TAFLP detects T-DNA insertion events by PCR amplification of genomic DNA flanking the T-DNA borders. Because the length and sequence of the flanking region depend on the genomic insertion site, independent transformation events are expected to yield distinct amplification patterns. In practice, the key point in assay design is to use a T-DNA border-specific primer in combination with an adaptor– or restriction-site-based primer so that only flanking fragments are amplified. The resulting PCR products are then visualized on agarose gels.

To confirm reproducibility, we analyzed four BASTA-resistant offspring from each of four previously generated independent transgenic lines. The assay yielded identical patterns among siblings, but clearly distinct patterns among independent lines (Fig. 2B). TAFLP can be applied to both the left and right T-DNA borders, and the two assays can complement each other (Fig. 2B). In addition, the flanking sequences can be identified by subsequent cloning and sequencing (Fig. 2A).

However, TAFLP has a detection limit of approximately 3 kb for PCR amplicon size, because fragments larger than 3 kb were rarely detected under our conditions. Amplicon-free patterns, which resemble the wild-type control, may occur when the adjacent restriction site is too distant from the T-DNA insertion site or when the T-DNA is inserted into a genomic region that is difficult to amplify by PCR, such as a highly heterochromatic region. Another limitation is that two independent insertion events may occasionally produce amplicons of similar length and therefore be indistinguishable with the gel system used. In our hands, TAFLP allowed screening of 384 plantlets in 5 working days, with a total hands-on time of 12 h, and the assay cost was only 0.2–0.3 euros per plantlet for one T-DNA border (Supplementary Table).

A total of 729 plantlets from the 10 transformation experiments described above was analyzed by TAFLP. On average, 16.2 plantlets shared the same TAFLP pattern, although the number ranged from 1 to 82 (Table 1; Fig. 2C; Supplementary Fig. 5). Notably, 17 T-DNA events were represented by fewer than five plantlets, indicating that the optimized tissue-culture procedure was effective in recovering minor or rare events. To compare the number of independent T-DNA events among experiments, the observed number of distinct TAFLP patterns was normalized to the number of embryos inoculated and expressed as the expected number of events per 100 embryos. These values ranged from 1.05 to 10.00, and most experiments yielded 1–3 events, although some reached 9–10 events (Table 1).

### Silwet L-77 potentially increases the number of initial inoculation events

TAFLP analysis suggested that the main limiting factor for recovering independent transgene insertion events was the number of initial inoculation events, since callus induction, selection, and regeneration had already been optimized and numerous plantlets derived from the same transgene event were often recovered. The semi-submergence strategy during the co-culture step described above was one attempt to increase the number of initial inoculation events (Fig. 1A,B), but further improvement was still possible.

Silwet L-77 (Silwet) is an organosilicone surfactant widely used in floral dip transformation of *Arabidopsis thaliana* (Clough and Bent, 1998) and other plant species (Curtis and Nam, 2001; Liu et al., 2008). We therefore hypothesized that Silwet treatment might also increase the number of initial inoculation events in maize embryos. In our first trial, Silwet was added to the co-culture medium at a final concentration of 0.01%; however, no embryos survived after 3 days of co-culture, whereas the control without Silwet induced callus normally, indicating that continuous exposure to Silwet during co-culture was detrimental to embryo viability (Fig. 3A).

**Figure 3.**
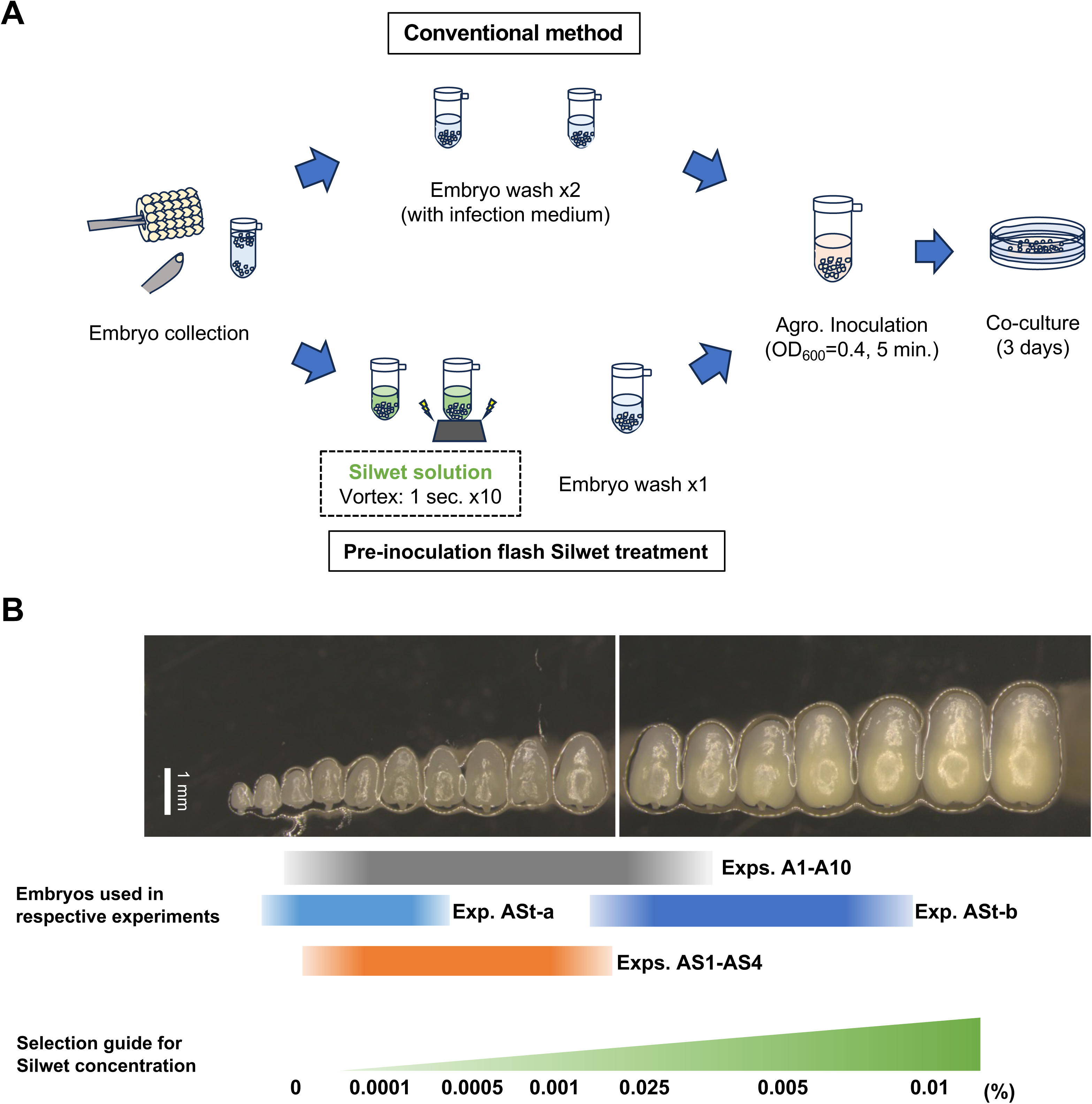
Application of Silwet treatment to potentially increase initial inoculation events. A. Schematic representation of the pre-inoculation flash Silwet treatment during the inoculation procedure for immature embryos. B. Size distribution of A188 embryos used in this study and a selection guide for Silwet concentration according to embryo size and morphology.

We therefore tested a brief Silwet treatment during the embryo-washing step, immediately after embryo isolation (Fig. 3A). Two independent embryo populations (ASt-a and ASt-b) were isolated and divided into three subpopulations, which were treated with 0, 0.0001, or 0.01% Silwet in the infection medium during the first washing step and then inoculated with the same Agrobacterium suspension carrying the *UBIpro:GUS* construct to visualize transformation events. One population (ASt-a) consisted of relatively small embryos, whereas the other (ASt-b) consisted of larger embryos that had not previously been used for inoculation (Fig. 3B). Inoculation events were visualized by X-Gluc staining after 3 days of co-culture (Supplementary Fig. 6A), and the remaining material was then processed through callus induction, selection, and regeneration as described above.

As shown in Table 1, in the ASt-a experiment with smaller embryos, 0.0001% Silwet treatment (ASt-a-2) resulted in 175.0% transformation efficiency (5.00 events per 100 embryos), whereas the 0% control (ASt-a-1) resulted in 46.2% efficiency (7.69 events per 100 embryos). In contrast, the 0.01% Silwet treatment (ASt-a-3) yielded no regenerants. In the ASt-b experiment with larger embryos, 0.0001% (ASt-b-2) and 0.01% (ASt-b-3) Silwet treatment resulted in 40.6% and 163.2% transformation efficiency, corresponding to 1.45 and 5.88 events per 100 embryos, respectively, whereas no plantlets were obtained from the 0% control (ASt-b-1; Supplementary Fig. 6B).

Finally, we performed four independent transformation experiments using flash treatment with 0.0001% Silwet and various constructs (Supplementary Table 1), which resulted in transformation efficiencies of 125.9%, 189.6%, 480.0%, and 671.4% (4.71, 6.94, 5.00, and 10.70 events per 100 embryos, respectively; Table 1 and Supplementary Fig. 7).

These results suggest that Silwet has a dual effect on initial inoculation events. First, flash Silwet treatment before embryo inoculation can increase transformation efficiency and is also applicable to larger embryos, which are otherwise difficult to transform using conventional methods. Second, there is a concentration window that depends on embryo size; excessive Silwet concentrations are harmful to smaller embryos and likely reduce transformation efficiency. Figure 3B provides a practical guide for selecting Silwet concentration according to embryo size and morphology, based on repeated experiments.

### Boosting method is applicable to the B104 inbred line

B104 is one of the most widely used inbred lines for Agrobacterium-mediated transformation in maize due to its amenability to transformation and shared origin with the maize reference genome line B73 (Raji et al., 2018; Aesaert et al., 2022). We therefore tested whether the optimized transformation method established here was also applicable to B104. In total, eight independent experiments were performed with various CRISPR/Cas9 constructs, resulting in transformation efficiencies ranging from 20.0% to 191.5% and an estimated number of events ranging from 3.08 to 9.57 per 100 embryos (Table 1; Supplementary Fig. 8).

This efficiency, which was comparable to that obtained with A188, was achieved using essentially the same tissue-culture procedure. However, two B104-specific features should be noted: (1) embryogenic callus tended to arise from the basal side of the embryo scutellum, closer to the suspensor, compared with A188 (Supplementary Fig. 9A); and (2) the embryo-size window suitable for transformation appeared narrower than in A188 (Supplementary Fig. 9B). Therefore, accurate timing of immature embryo harvesting was likely more important for B104. Although the effect of Silwet treatment was not tested in this line, CoReCa also showed a morphology similar to that observed in A188 (Supplementary Fig. 9C). We further performed six tracing experiments starting from single CoReCa after Sel-3 and found that each CoReCa produced 30, 56, 8, 10, 14, and 14 plantlets, respectively. TAFLP analysis of these plantlets suggested that they mostly shared the same T-DNA event; however, in two cases, two independent events were isolated from a single CoReCa, indicating that these calli were chimeric (Supplementary Fig. 8).

Overall, these results demonstrate that the optimized transformation platform can be successfully transferred to another widely used maize inbred line, although some genotype-specific adjustments are required.

### Boosting method is also applicable to other selection systems: hygromycin and G418

Next, we examined whether the method was applicable to selection systems other than BASTA. For this purpose, we constructed *UBIpro:GUS* vectors carrying either *hygromycin phosphotransferase II* (*hptII*) or *neomycin phosphotransferase II* (*nptII*), driven by the *CaMV 35S* promoter, for hygromycin B or G418 selection, respectively, and transformed them into the A188 inbred line. Selection pressure was increased gradually across the selection phases, following the scheme used for the BASTA system: 15, 30, and 40 mg/L hygromycin B, and 10, 20, and 40 mg/L G418 in Sel-1, Sel-2, and Sel-3 media, respectively.

In these initial trials, we recovered 24 and 19 plantlets under hygromycin B and G418 selection, corresponding to transformation efficiencies of 27.0% and 15.4%, respectively. In both cases, the recovered plantlets originated from a single event (1.12 and 0.96 events per 100 embryos, respectively; Table 1). GUS expression in the regenerating plantlets confirmed successful transgene integration (Supplementary Fig. 10A), and transmission to the T1 progeny confirmed the heritability of the events (Supplementary Fig. 10B), consistent with the results obtained using the BASTA selection system. One notable difference from BASTA selection was that CoReCa formation was less distinct after the Sel-3 phase when using the hygromycin and G418 systems. Instead, clear selection outcomes were observed only after Reg-1, suggesting that maize callus responds differently to different selective agents.

We also tested whether hygromycin and G418 selection systems were suitable for CRISPR/Cas9 mutagenesis. To this end, we generated three constructs carrying *bar*, *hptII*, or *nptII* selection markers and sharing the same CRISPR/Cas9 cassette targeting the maize *MULTIPLE ARCHESPORIAL CELLS 1* (*MAC1*) gene (Fig. 4A) (Wang et al., 2012). We then performed transformation experiments in A188 (designated A11, AH2, and AG2, respectively; Table 1 and Supplementary Table 3). This resulted in 39, 130, and 90 plantlets, corresponding to transformation efficiencies of 19.7%, 191.2%, and 50.3% under BASTA, hygromycin, and G418 selection, respectively. The expected number of events was 2.9 and 2.2 per 100 embryos for the hygromycin and G418 systems, whereas this value was not determined for the BASTA system in this experiment.

**Figure 4.**
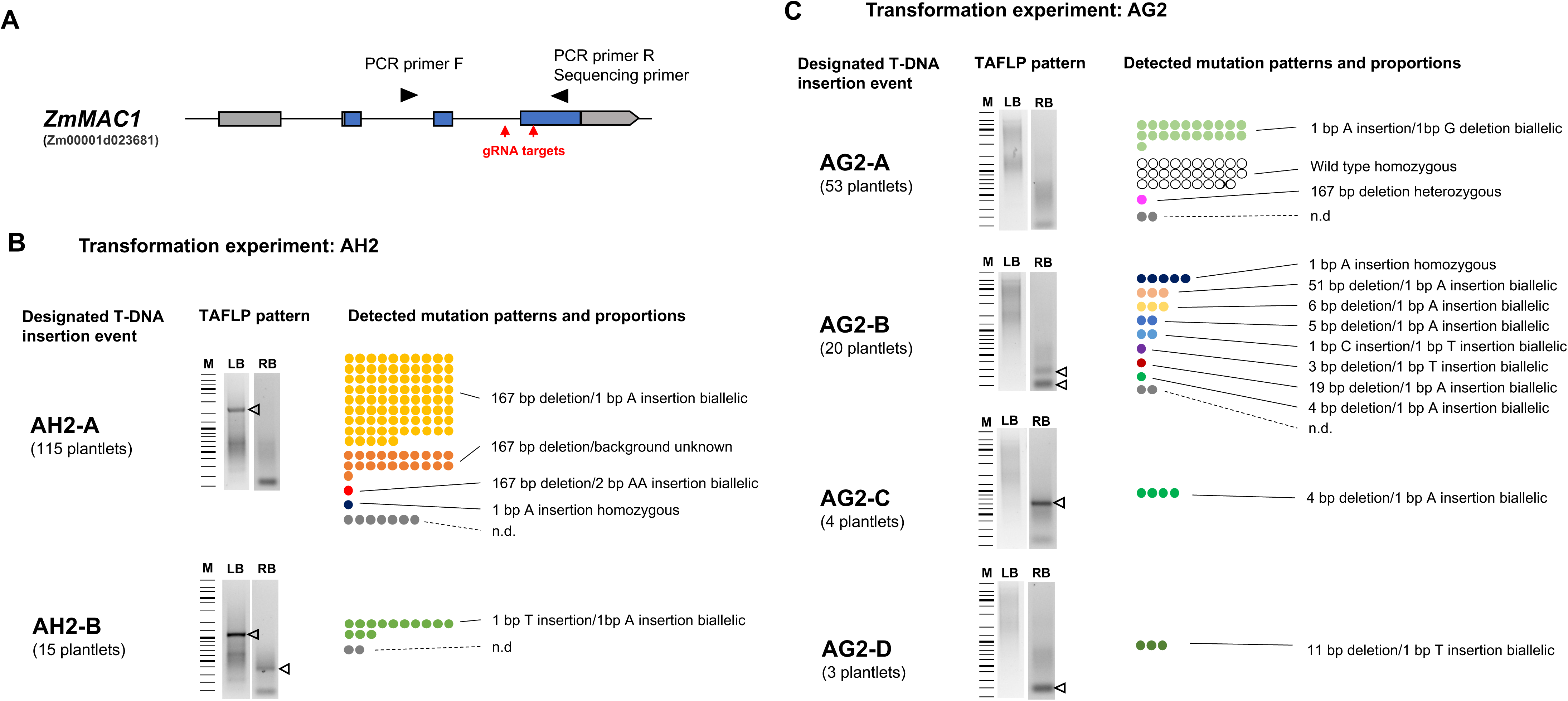
Regeneration through splitting is highly suitable for isolating independent and minor target-editing events with CRISPR/CasG. A. Schematic representation of the MAC1 target locus showing guide RNA (gRNA) target sites and PCR/sequencing primer positions. B. Proportions of detected mutation events in respective T-DNA insertion events determined by TAFLP in transformation experiment AH2. Each dot represents one plantlet and is color-coded according to mutation event. n.d., mutation event not determined due to PCR or sequencing failure. TAFLP patterns are shown using the same scheme as in Fig. 2C. C. Proportions of detected mutation events in transformation experiment AG2, shown using the same scheme as in Fig. 4B.

We then screened all recovered plantlets for mutations at the *MAC1* target site by PCR amplification of the target region followed by Sanger sequencing. In total, 94.9% and 94.6% of plantlets carried mutations in *MAC1* under BASTA and hygromycin selection, respectively, whereas 62.2% of plantlets recovered under G418 selection were mutated. In most cases, the mutations were biallelic (see below). Together, these results indicate that CRISPR/Cas9 mutagenesis is highly effective across all three selection systems.

### Regeneration through splitting is highly beneficial for CRISPR/CasG mutagenesis

For CRISPR/Cas9 mutagenesis, once the construct is integrated into the genome, Cas9 can generate independent editing events in different cells during regeneration. Therefore, even plantlets derived from the same T-DNA insertion event may carry different target-site mutations. The regeneration-through-splitting strategy is particularly useful in this context because it increases the chance of recovering distinct editing outcomes from a shared transgenic event. To evaluate this, we analyzed mutation profiles at the *MAC1* locus together with T-DNA event profiles determined by TAFLP.

In experiment AH2, two clearly distinguishable transformation events were detected (Fig. 4B). In the major subpopulation carrying T-DNA event AH2-A, most plantlets (106 of 115) carried a 167-bp deletion in at least one allele, whereas the second allele often contained a 1-bp A insertion, although in some cases the sequence was unclear, likely because of PCR bias at the target region (Fig. 4B). Two plantlets showed mutation patterns clearly different from the others: one carried a 167-bp deletion/2-bp AA insertion as a biallelic genotype, and the other was homozygous for a 1-bp A insertion. The minor subpopulation carrying T-DNA event AH2-B predominantly showed 1-bp T and A insertions as biallelic mutations (13 of 15 plantlets; Fig. 4B).

In experiment AG2, four distinguishable TAFLP patterns were detected (Fig. 4C). The minor T-DNA events AG2-C (4 plantlets) and AG2-D (3 plantlets) were characterized by a 4-bp deletion/1-bp T insertion and an 11-bp deletion/1-bp A insertion, respectively, as biallelic mutations (Fig. 4C). The second major subpopulation carrying T-DNA event AG2-B was highly variable, with eight distinct allelic combinations identified among 18 plantlets (Fig. 4C). The largest subpopulation carrying T-DNA event AG2-A consisted of 21 plantlets with a 1-bp T insertion/1-bp C deletion as a biallelic genotype, 29 plantlets with wild-type sequences, one plantlet with a heterozygous 167-bp deletion, and two plantlets with unclear sequencing results due to PCR limitations (Fig. 4C). Notably, AG2-A may have represented a mixture of multiple T-DNA events, because TAFLP patterns specific to this subpopulation were not detected with either left– or right-border assays, suggesting that these events were beyond the detection limit of TAFLP.

Together, these two case studies lead to two main conclusions. First, recovering a higher number of distinct T-DNA insertion events is advantageous, because different T-DNA events tended to produce different mutation types and combinations, reflecting differences in Cas9 activity. Second, even among plantlets derived from the same T-DNA event, regeneration through splitting was effective for recovering rare or unique mutations at the target locus. Overall, our method is well suited for CRISPR/Cas9-mediated mutagenesis and additionally allows recovery of minor T-DNA events (Fig. 2C; Supplementary Fig. 5).

### Case study 1: Isolation of new *mac1* mutant lines

Although CRISPR/Cas9 has advanced rapidly, it can still be challenging to isolate and establish mutant lines for genes that are essential for reproduction. Using our method, we isolated mutants of two previously reported meiosis-related maize genes, for which loss of function causes severe sterility.

The first case targeted *MULTIPLE ARCHESPORIAL CELLS 1* (*MAC1*), a gene essential for anther development and microsporocyte specification (Sheridan et al., 1996; Wang et al., 2012). In experiment A11, using a BASTA-selection CRISPR/Cas9 construct targeting *MAC1*, we recovered 39 T0 plantlets, all of which shared the same mutation pattern: a 197-bp deletion/1-bp A insertion as a biallelic genotype (Supplementary Fig. 11). All T0 plantlets were completely male sterile, as previously reported (Wang et al., 2012). However, previous studies showed that *mac1-1* mutants exhibit strong female sterility but can produce viable seeds at low frequency (Sheridan et al., 1996). We therefore performed extensive backcrossing with wild-type pollen using the cobs of these male-sterile T0 plants, followed by embryo rescue. This allowed us to recover six T1 plants, five of which carried the CRISPR/Cas9 transgene and either the 197-bp deletion or the 1-bp A insertion at the *MAC1* locus, consistent with segregation from the parental T0 plant (Supplementary Fig. 11).

Three of these CRISPR/Cas9-positive T1 plants showed complete male sterility, indicating that the transgene remained active in the T1 generation and generated de novo mutations in the remaining wild-type *MAC1* allele. In contrast, two CRISPR/Cas9-positive T1 plants, both carrying the 197-bp deletion, showed a chimeric phenotype with fertile and sterile sectors within a single tassel. This enabled recovery of T2 seeds by backcrossing using viable pollen from the fertile sectors. Among the T2 seeds, we successfully isolated heterozygous plants with the 197-bp deletion that were free of the CRISPR/Cas9 construct, and we subsequently established the *mac1-2* segregating population by self-crossing (Supplementary Figs. 11 and 12A). Similarly, from one fertile T1 plant carrying the 1-bp A insertion (designated *mac1-3*; Supplementary Fig. 12A), we established a CRISPR/Cas9-free segregating population by self-crossing.

The phenotypes of *mac1-2* and *mac1-3* were examined in the segregating populations. Both mutants showed vegetative growth comparable to that of wild-type siblings (Supplementary Fig. 12B), but exhibited complete male sterility with unopened tassel spikelets and shriveled anthers lacking viable pollen, as well as reduced female fertility in crosses with wild-type pollen (Supplementary Fig. 12C,D). Confocal laser-scanning microscopy of anthers at meiotic prophase I showed that, in both mutants, the inner anther wall layers (tapetum, middle layer, and endothecium) were abnormally developed and not properly differentiated, whereas these layers were clearly distinguishable in wild-type siblings (Supplementary Fig. 12E,F). These phenotypes were consistent with the previously reported *mac1-1* phenotype (Sheridan et al., 1996; Wang et al., 2012), indicating that we successfully isolated new loss-of-function alleles of *MAC1* in the A188 inbred line.

### Case study 2: Isolation of *acoz1-3* and characterization using reporter lines

The second case study focused on the maize *ABNORMAL CHROMOSOME ORGANIZATION IN ZYGOTENE 1* (*ACOZ1*) gene, which encodes an F-box protein essential for chromosome behavior and organization during meiosis (Jing et al., 2022). Loss-of-function mutants exhibit complete male and female sterility. To isolate a new *acoz1* allele in this study, we designed two guide RNAs targeting the first and third introns of *ACOZ1*, with the aim of generating a large deletion spanning the intervening region rather than small indels at the individual target sites (Fig. 5A; Supplementary Fig. 13A).

**Figure 5.**
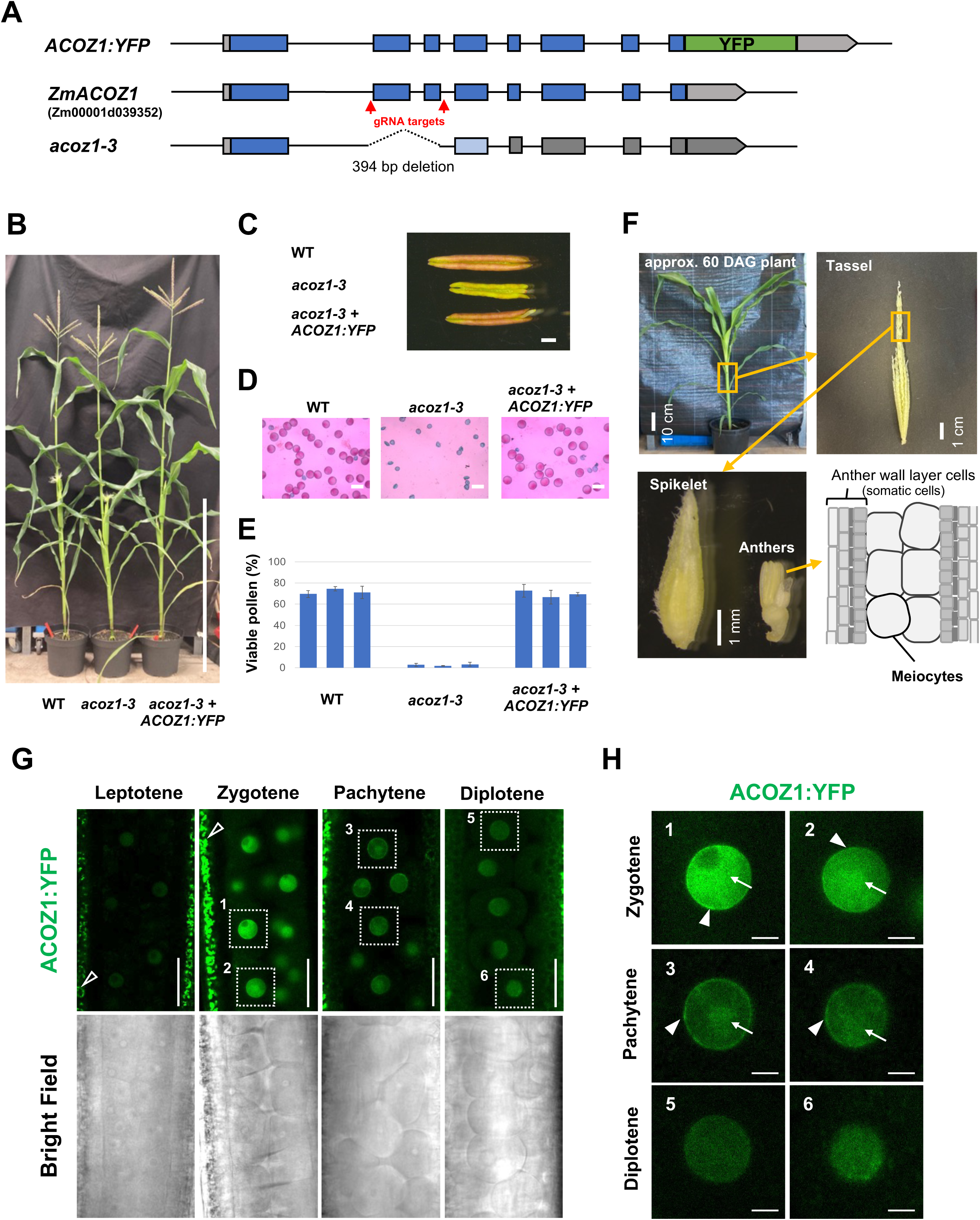
Phenotypic complementation of the acoz1-3 mutant by an ACOZ1:YFP reporter. A. Structures of the *ACOZ1* locus showing guide RNA (gRNA) target sites, the isolated *acoz1-3* mutant allele, and the ACOZ1:YFP reporter construct. B. Plant morphology of wild-type (WT), *acoz1-3*, and a complemented line carrying ACOZ1:YFP in the *acoz1-3* background at the flowering stage. Bar = 1 m. C. Anther morphology at the flowering stage. Bar = 1 mm. D. Pollen fertility assessed by Peterson’s staining. Bars = 100 µm. E. Quantification of pollen fertility. Each bar represents the mean of individual plants determined from three independent anthers, and error bars indicate standard deviation. F. Plant, tassel, and spikelet morphology at the meiotic stage. Bottom right: longitudinal section of a meiotic anther observed by confocal microscopy. G. Accumulation pattern of ACOZ1:YFP during prophase I. Images were acquired and processed using identical microscopy settings across samples. Open arrowheads indicate chlorophyll autofluorescence from endothecium cells. Bars = 50 µm. H. Subcellular localization of ACOZ1:YFP during prophase I. Numbers correspond to the insets shown in Fig. 5G. Arrows and arrowheads indicate chromosomal and nuclear peripheral localization of ACOZ1:YFP, respectively. Bars = 10 µm.

From the transformation experiments with the *ACOZ1*-targeting CRISPR/Cas9 construct, we recovered 284 plantlets and screened them by PCR for putative deletion events. Fourteen plantlets showed substantially shorter PCR fragments than the wild-type allele, and 13 of these were successfully backcrossed to wild-type plants (Supplementary Fig. 13A,B). From one T1 seed population derived from these crosses, we recovered two CRISPR/Cas9-negative plants that were heterozygous for a 394-bp deletion allele of *ACOZ1* and lacked the entire second and third exons, corresponding to the F-box coding region. This allele was designated *acoz1-3* (Fig. 5A). The *acoz1-3* segregating T2 population was maintained by subsequent sib-crossing (Supplementary Fig. 13B). As expected, *acoz1-3* mutants showed vegetative growth comparable to that of wild-type siblings but severe male and female sterility, consistent with previous reports (Supplementary Fig. 14) (Jing et al., 2022).

In parallel, we generated *ACOZ1:YFP* reporter lines to visualize ACOZ1 expression during meiosis. Two independent events, derived from separate transformation experiments, were screened by confocal laser-scanning microscopy and maintained by backcrossing with wild-type plants. A T1 plant from one event, showing higher expression and a single-locus segregation pattern, was crossed with the *acoz1-3* heterozygous line and subsequently self-crossed to obtain an F2 (T3) line for complementation analysis (Supplementary Fig. 13B). Plants carrying the *ACOZ1:YFP* reporter in the *acoz1-3* mutant background showed a fertile phenotype comparable to that of wild-type siblings (Fig. 5B–E), indicating that the introduced *ACOZ1:YFP* construct was functional.

ACOZ1:YFP was detected in meiocyte nuclei throughout prophase I, with peak expression at zygotene (Fig. 5F,G). At zygotene and pachytene, ACOZ1:YFP showed a chromatin-like localization pattern (Fig. 5H, arrows) and was also enriched at the nuclear periphery (Fig. 5H, arrowheads), consistent with a possible role in meiotic chromosome organization and localization at the nuclear envelope region.

In rice, the inner nuclear-envelope-localizing SUN-domain protein OsSAD1 showed an eccentric localization pattern in wild-type meiocytes at zygotene, whereas this pattern was absent in the *zygotene 1*(*zygo1*) mutant, the rice ortholog of maize *ACOZ1*(Zhang et al., 2017). To test whether maize *acoz1-3* mutants show a similar change in SUN protein localization, we generated a maize *SUN1:RFP* reporter and introduced it into the *acoz1-3* background by crossing, followed by confocal microscopy (Supplementary Fig. 15A–C). In *acoz1-3* heterozygous siblings used as controls, 69.8% of meiocytes showed eccentric SUN1:RFP localization at the nuclear equator, which we defined as the “crescent” pattern (Supplementary Fig. 15C,D). In contrast, in the *acoz1-3* mutant, 74.6% of zygotene-like cells showed signal decorating the entire nuclear equator circumference, a configuration of SUN1:RFP defined as the “full moon” pattern (Supplementary Fig. 15C,D). These observations indicate that ACOZ1 is required for the eccentric localization pattern of SUN1 in maize meiocytes at zygotene.

It was recently reported that nuclear lamina proteins are rapidly and specifically removed from early prophase I meiocytes in *Arabidopsis*, and that this process is regulated by an SCF complex involving the F-box proteins REDUCED MALE FERTILITY 1/2 (RMF1/2), which are orthologous to maize ACOZ1 (Yuan et al., 2025). To test whether ACOZ1-dependent degradation of maize nuclear lamina proteins also occurs during meiosis, we generated reporter constructs for maize NMCP/CRWN homologs 1 and 2 (NCH1 and NCH2; Gumber et al., 2019), fused to RFP at the C terminus, and introduced them into maize plants. In these experiments, transformation was performed using embryo populations derived from crosses between established *acoz1-3* heterozygous plants and wild-type plants, thereby bypassing time-consuming subsequent crosses (Supplementary Fig. 16).

Using TAFLP, *acoz1-3* genotyping, and confocal screening, we recovered two independent events for both *NCH1:RFP* and *NCH2:RFP* in the *acoz1-3* heterozygous background, and crossed them with *acoz1-3* heterozygous plants. This allowed us to assess *NCH:RFP* expression in *acoz1-3* segregating populations already at the T1 generation (Supplementary Fig. 16). In wild-type siblings, both NCH1:RFP and NCH2:RFP were clearly observed at the nuclear periphery in premeiotic stages but were barely detectable in meiocytes at leptotene, zygotene, and early pachytene, although signals were clearly detected in somatic anther wall cells at these stages (Fig. 6A,B; Supplementary Fig. 17A,B). NCH:RFP signals reappeared in meiocytes at the later diplotene stage (Fig. 6A,B; Supplementary Fig. 17A,B). In contrast, in all *acoz1-3* mutant cases, clear NCH:RFP signals were detected in nuclei of zygotene– and pachytene-like meiocytes, indicating that NCH degradation did not occur in *acoz1-3* mutant meiocytes (Fig. 6A,B; Supplementary Fig. 17A,B).

**Figure 6.**
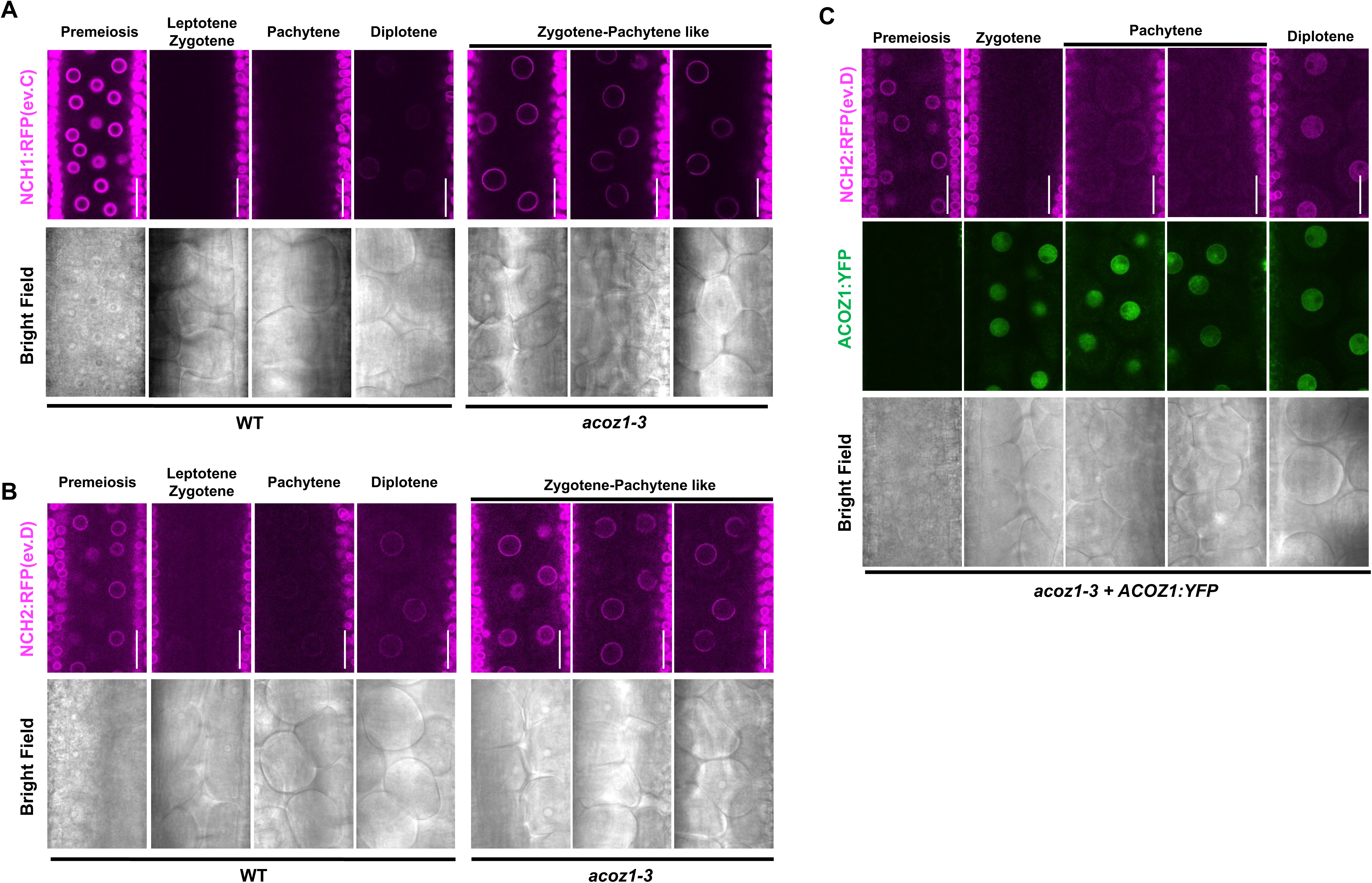
Nuclear lamina proteins NCH1 and NCH2 are targeted and degraded by ACOZ1 during early prophase I. A. Accumulation pattern of NCH1:RFP (ev.C, independent event C) in meiotic anthers compared with wild-type and *acoz1-3* siblings. Bars = 50 µm. B. Accumulation pattern of NCH2:RFP (ev.D) in meiotic anthers compared with wild-type and *acoz1-3* siblings. Bars = 50 µm. C. Accumulation pattern of NCH2:RFP (ev.D) in meiotic anthers of the complemented line carrying ACOZ1:YFP in the *acoz1-3* mutant background. Bars = 50 µm.

Finally, expression of one NCH2:RFP event (designated ev.D) was examined in a complemented *acoz1-3* background carrying *ACOZ1:YFP*. In this background, NCH2:RFP was again barely detectable during early prophase I, and its expression pattern was nearly opposite to that of ACOZ1:YFP, suggesting that functional ACOZ1 targets NCH proteins for degradation in maize meiocytes during early prophase I (Fig. 6C).

## Discussion

Although the mechanisms of Agrobacterium-mediated transformation has been understood in detail and its application is now routine in several model species (Singh and Prasad, 2016; Gelvin, 2010), maize continues to be limited by genotype dependence, low recovery of independent events, and strong sensitivity to tissue culture conditions (Rahnama, 2010; Singh and Prasad, 2016). Our work shows that transformation performance can be substantially improved by reconsidering tissue culture not as a fixed background step, but as a series of developmental and physiological transitions that can each be tuned to favor infection, embryogenic competence, selection, and regeneration.

A key implication of this study is that successful transformation depends on matching the timing and position of infection with the developmental origin of the regenerative tissue (Fig. 1B, Supplementary Figs. 2 and 9). Rather than treating embryo inoculation, callus induction, and regeneration as independent steps, our data suggest that these phases must be spatially and temporally coordinated to increase the likelihood that transformed cells contribute to regenerable tissue (Fig. 1A). This framework helps explain why relatively simple changes, such as embryo submergence, resting phases, and controlled embedding, can have strong effects when combined appropriately. In this sense, the method illustrates how transformation efficiency is shaped by tissue architecture and developmental dynamics as much as by bacterial delivery itself. We further postulate that this framework can be applied to the transformation procedures of other species, which have currently low transformation rates such as Sorghum and wheat (Li et al., 2012; HAN and CAI, 2024).

Another important observation is that the Silwet treatment used during inoculation appears to contribute to improved transformation in maize in a manner that differs from its well-known role in the Arabidopsis floral dip method. In Arabidopsis, surfactants are thought to enhance Agrobacterium access to target gametophytic cells hidden within floral tissues (Clough and Bent, 1998). In maize immature embryos, however, the relevant target cells are the scutellum epidermis, a single-cell layer that is already exposed on the embryo surface and gives rise to embryogenic callus during somatic embryogenesis (Springer et al., 1979; Vasil et al., 1985). Thus, while Silwet may still modify the local microenvironment and improve bacterial contact with epidermal cells, its effect in maize is unlikely to depend primarily on increasing tissue accessibility in the same way as in floral dip.

Instead, a more plausible explanation is that Silwet imposes a mild stress or wounding-like condition that alters the competence of scutellum cells for somatic embryogenesis. Wounding is known to trigger callus formation and dynamic changes in gene expression involving key developmental regulators (Iwase et al., 2021), and Silwet treatment may mimic some of these responses. This interpretation is consistent with our empirical observation that well-developed immature embryos did not always yield the highest transformation efficiencies, even when they fell within the acceptable size range. In contrast, embryos derived from stressed parent plants, such as those exposed to heat or cold stress, or embryos with slightly abnormal morphology, sometimes performed better.

Similarly, Ishida et al. (2007) reported that heat shock prior to Agrobacterium infection can improve transformation, suggesting that external stress-inducing treatments may increase the competency of scutellum cells. In this context, the Silwet treatment developed here may provide a flexible and fine-tunable way to adjust embryo responsiveness depending on genotype, growth condition, or embryo quality (Fig. 3). We also postulate that identifying a critical Silwet concentration that possibly induces a mild stress could boost transformation of other species, too.

The frequent callus-splitting strategy, referred to here as regeneration through splitting, likely exploits another underappreciated principle: regeneration is favored by maintaining tissues in small, actively responsive units rather than in large, heterogeneous masses. This may reflect improved access to nutrients, hormones, and oxygen, or the ability of smaller callus fragments to retain higher regenerative competence (Fig. 1A,D). Whatever the precise mechanism, the approach appears particularly advantageous when the goal is not only to recover transformants, but also to increase the number of distinct genetic events available from a transformed population (Figs. 2C and 4; Supplementary Figs. 5–8). This is especially relevant for genome editing, where the biological value of a transformation experiment depends on the diversity of mutation outcomes recovered.

The value of recovering multiple plantlets is not limited to distinct events. Even among plants sharing the same T-DNA insertion or the same editing outcome, the availability of several independent regenerants can be useful as biological backups depending on downstream objectives. For example, in the isolation of the *mac1* new allele, extensive backcrossing among multiple plantlets carrying the same sterile biallelic mutation made it possible to obtain stable mutant lines (Supplementary Figs. 11 and 12). Similarly, the ability to distribute favorable insertion events, such as high-expressing reporter constructs, or desired mutant alleles across multiple crossing partners may accelerate the generation of reporter lines or higher-order mutants, thereby supporting functional genomic studies in maize. This is particularly important because regenerated T0 plantlets often show disadvantageous reproductive morphology, including reduced cob and tassel size, and these effects can be more severe after prolonged tissue culture. From this perspective, obtaining both multiple plantlets with the same event and multiple distinct events is highly beneficial for maize functional genomics.

TAFLP provides a practical complement to this strategy by helping distinguish independent insertion events from redundant regenerants. Although not intended as a definitive mapping method, it offers a rapid and inexpensive way to estimate event complexity early in the pipeline, which is valuable when large numbers of regenerants are produced. Together, regeneration-through-splitting and TAFLP support a more event-aware transformation workflow that should reduce redundancy and improve downstream experimental efficiency. For reporter constructs in particular, the insertion site is important because transgene expression is strongly influenced by the surrounding chromatin landscape, including whether the locus is heterochromatic or euchromatic. TAFLP therefore provides a useful first overview to guide more targeted screening of transgene activity, such as microscopy or RNA– or protein-based expression analyses. By contrast, TAFLP may be less essential for CRISPR/Cas9 mutagenesis, where the structure of the editing outcome at the target site is often more relevant than insertion complexity. Choosing screening strategies according to the experimental purpose should further increase the practical value of the method described here.

The broad applicability of the platform across different maize genotypes, selection systems, and CRISPR/Cas9 applications suggests that the approach captures general features of maize tissue culture rather than a narrow genotype-specific optimization. This is encouraging because one persistent obstacle in maize biotechnology is the gap between protocols that work well in principle and those that remain robust across users, constructs, and experimental goals. By focusing on the tissue culture phase itself, our study provides a framework that may be transferable to other recalcitrant crop species where transformation is constrained by similar developmental bottlenecks.

More broadly, our results support the idea that improving plant transformation will increasingly depend on integrating developmental biology with biotechnology. The most effective strategies may not be those that simply intensify selection or increase bacterial exposure, but those that better align transformation conditions with the biology of the target tissue. In this respect, tissue culture optimization is not merely technical refinement; it is an opportunity to reshape the cellular context in which transgenesis and genome editing occur.

In summary, this work provides a practical transformation framework and highlights principles that may be useful beyond maize. A more explicit consideration of tissue state, regeneration dynamics, and event structure should help improve transformation workflows for functional genomics and crop engineering more generally.

## Materials and Methods

### Plant materials and growth conditions

Wild-type maize inbred lines A188 and B104, as well as seed-derived generations of transgenic lines, were germinated in soil trays with cells measuring 8 cm in diameter and 6 cm in height, filled with CL-ED73 soil (Werner Tantau GmbH), and grown in a closed greenhouse chamber at the Institute of Plant Science and Microbiology, University of Hamburg. Plants were maintained at a constant temperature of 26°C under natural light supplemented with growth lamps to provide a 14/10 h light/dark photoperiod. Plants at approximately 28 days after germination (DAG) were transferred to larger pots (24 cm diameter, 22 cm height) filled with CL-ED73 soil supplemented with 40 g Hauert Long fertilizer (Hauert MANNA Düngerwerke GmbH). Transgenic T0 plantlets were grown under the same conditions, except that the first 2 weeks in cell trays were used for acclimatization.

### Vector construction and general setup for Agrobacterium-mediated transformation

Binary vector backbones p7oM-LH, p6oM-LH, and p9oM-LH (DNA Cloning Service, https://dna-cloning.com) were used for BASTA (bar), hygromycin (hptII), and G418 (nptII) selection, respectively. Detailed construct structures are provided in Supplementary Table 1. For reporter constructs, the binary vectors were first modified for Gateway® cloning by introducing an attR1-CmR-ccdB-attR2 cassette. YFP-, mRFP-, or GUS-fused reporter genes cloned into pENTR2B entry vectors were transferred into the binary vectors using LR Clonase II enzyme mix (Invitrogen).

For CRISPR/Cas9 constructs, three Cas9 expression modules were generated: one expressing SpCas9 (Char et al., 2017) or SaCas9 (Fauser et al., 2014) under the maize *UBIQUITIN1* (*UB1*) promoter, and one expressing SpCas9 under the maize *DISRUPTED MEIOTIC cDNA1*(*DMC1*) promoter (Feng et al., 2018) combined with a Ubiquitin1 promoter-driven RUBY marker (He et al., 2020). Guide RNA expression entry vectors for SpCas9 were obtained from Char et al. (2017), and those for SaCas9 were modified accordingly. Guide RNA sequences were cloned as previously described (Char et al., 2017) and combined with the compatible Cas9 expression binary vectors via LR cloning.

Constructed binary vectors were introduced into *Agrobacterium tumefaciens* strain LBA4404, which harbored additional *virB* and *virG* genes from *pTiBo542* (Komari et al., 1986), by electroporation. Immature embryos were collected at 10–13 days after pollination (DAP) from A188 and 11–14 DAP from B104; the exact timing varied with the growing season. Embryos were subjected to *Agrobacterium* inoculation. The detailed transformation and tissue culture procedures are provided in Supplementary Procedure 1.

Histochemical GUS staining (Fig. 1B, Supplementary Fig. 6A, and Supplementary Fig. 10) was performed according to Brettschneider et al. (1997).

## Genetic analyses of maize plants

In some cases, T1 or later generations of BASTA-selected lines were tested for transgene integration by spraying seedlings at 10 DAG with 200 mg/L BASTA solution (Bayer Crop Science) containing 0.1% Tween-20. In other cases, PCR genotyping was performed to assess the presence or absence of transgenes or to detect mutations in genes of interest (primer sequences are listed in Supplementary Table 2). Genomic DNA was prepared as previously described with minor modifications (Dellaporta et al., 1983). For TAFLP analysis, DNA samples were additionally treated with 3 µg RNase A (New England Biolabs) and purified by ethanol precipitation. The TAFLP procedure is described in Supplementary Procedure 2. For mutation analysis of the MAC1 locus (Fig. 4), the target region was amplified by PCR and purified using MagMAX™ Pure Bind Beads (Applied Biosystems), followed by Sanger sequencing at GENEWIZ Germany GmbH.

## Fertility analyses

Pollen fertility was examined according to Peterson et al. (Peterson et al., 2010). To avoid bias from uneven pollen distribution on the test slide, all pollen grains collected from a single anther in an unopened spike were examined, and three replicates were analyzed per plant. For the female fertility test, an adequate amount of freshly collected wild-type pollen was crossed to well-emerged silks on the cob (5–7 days after first silk emergence). This cross was repeated three times at 1-day intervals, and cobs were examined for seed set at 40 DAP.

## Confocal laser scanning microscopy

Anthers at meiotic stages were dissected from plants at approximately 60 DAG, placed on a slide with sterile water, and covered with a coverslip. Fluorescence and bright-field images were acquired using a Zeiss LSM900 or Zeiss LSM880 confocal microscope equipped with a C-Apochromat 40×/1.2 W objective and Zen 2.3 SP1 software (Carl Zeiss). Images were acquired using identical settings across comparable experiments and processed uniformly with Fiji software.

## Author contributions

S.O., Conceptualization, Investigation, Formal analysis, Writing; M.O., Investigation; R.B., Investigation; D.S., Investigation; K.M., Investigation; M.B., Investigation; M.v.d.H., Investigation; A.S., Conceptualization, Funding acquisition, Project administration, Resources, Supervision, Writing.

## Acknowledgments

This work was supported by the German Research Council (Deutsche Forschungsgemeinschaft, DFG) via Research Unit 5235 (CSCS: Cereal Stem Cell Systems) to A.S. (SCHN 736/20-1).

## Supplementary Material

**Supplementary Table 1. Summary of constructs generated and used in this study.**

**Supplementary Figure 1. Effect of meropenem on embryonic callus formation.**

**Supplementary Figure 2. Semi-submerging embryos during co-culture.**

**Supplementary Figure 3. Effect of embryo/callus embedding during embryonic callus formation and the early phase of selection.**

**Supplementary Figure 4. Glufosinate metabolism in the presence or absence of the bar marker gene and its effect on tissue culture conditions.**

**Supplementary Figure 5. Proportions of designated T-DNA insertion events in transformation experiments A1, A2, A3, A5, A7, A8, and A10.**

**Supplementary Figure 6. Test experiments for the application of Silwet during inoculation.**

**Supplementary Figure 7. Proportions of designated T-DNA insertion events in transformation experiments with Silwet treatment (AS1–AS4).**

**Supplementary Figure 8. Proportions of designated T-DNA insertion events in transformation experiments using the B104 inbred line (B1–B8).**

**Supplementary Figure 9. Immature embryo and callus morphology of the B104 inbred line.**

**Supplementary Figure 10. GUS histochemical staining of regenerating plantlets from transformation experiments with hptII (hygromycin) and nptII (G418) selection systems.**

**Supplementary Figure 11. Schematic representation of the crosses and screenings used to isolate mac1-2 and mac1-3 mutants.**

**Supplementary Figure 12. Characterization of the mac1-2 and mac1-3 mutant phenotypes.**

**Supplementary Figure 13. Schematic representation of the crosses and screenings used to isolate the acoz1-3 mutant, generate the ACOZ1:YFP reporter, and establish the acoz1-3 complementation line carrying ACOZ1:YFP.**

**Supplementary Figure 14. Characterization of the acoz1-3 mutant phenotype.**

**Supplementary Figure 15. Characterization of SUN1:RFP localization in acoz1-3 mutant meiocytes.**

**Supplementary Figure 16. Schematic representation of the transformation experiments, crosses, and screenings used to establish NCH:RFP reporters in the acoz1-3 mutant background.**

**Supplementary Figure 17. Additional examples of NCH:RFP reporter expression in the acoz1-3 mutant background.**

## Supplementary Figure Legends

**Supplementary Figure 1. Effect of meropenem on embryonic callus formation.**

Immature embryos after the Sel-I phase were cultured with 100 mg/L carbenicillin (left) or 25 mg/L meropenem (right). Embryos were derived from the same inoculation event. Bottom images show magnified views of the embryos in the upper insets. Arrowheads indicate successful formation of embryonic callus.

**Supplementary Figure 2. Semi-submerging embryos during co-culture**.

Immature embryos before (left) and after (right) the co-culture phase, showing optimal semi-submerging in Agrobacterium infection medium. After 3 days of co-culture, the embryos enlarged and the infection medium became cloudy due to bacterial growth, indicating successful co-culture. The illustration at the bottom shows the optimal degree of semi-submerging based on the structure of immature embryos from top and side views. Bars = 1 mm.

**Supplementary Figure 3. Effect of embryo/callus embedding during embryonic callus formation and the early phase of selection**.

Immature embryos after the Sel-I phase were cultured without embedding (left) or with embedding (right) during the resting and Sel-1 phases. Embedding was performed according to the guide shown in Fig. 1A. Bottom images show magnified views of the embryos in the upper insets. Note that the no-embedding experiment is identical to the meropenem-treated experiment shown in Supplementary Fig. 1. These two experiments were not derived from the same inoculation event, but from embryo samples of similar size and morphology. Arrowheads indicate successful formation of embryonic callus.

**Supplementary Figure 4. Glufosinate metabolism in the presence or absence of the bar gene and its effect on medium pH**.

A. Comparison of glufosinate metabolism in bar-positive cells (top) and wild-type cells (bottom). Note that wild-type cells release ammonium ions (NH4+) from glufosinate.

B. Comparison of pH changes in Reg1 medium before (left) and after 10 days of culture with non-resistant calli (middle) or with some resistant calli (right). Dashed circles indicate the locations of resistant calli. Medium pH was measured using pH-indicator paper (pH 1–14 Universal Indicator; Merck). The color chart for the respective pH range is shown at the bottom. The medium cultured with non-resistant calli showed pH 7.0–7.5, whereas the optimal pH of Reg1 medium is 5.8.

**Supplementary Figure 5. Proportions of designated T-DNA insertion events in transformation experiments A1, A2, A3, A5, A7, A8, and A10**.

The proportions of designated T-DNA insertion events in the respective transformation experiments are shown using the same structure as in Fig. 2C.

**Supplementary Figure 6. Test experiments for Silwet treatment prior to Agrobacterium inoculation**.

A. Histochemical GUS staining of inoculated embryos after the co-culture phase following treatment with various Silwet concentrations. Arrowheads indicate embryos showing GUS spots in the potential callus-inducing area.

B. The proportions of designated T-DNA insertion events in the respective transformation experiments are shown using the same structure as in Fig. 2C.

**Supplementary Figure 7. Proportions of designated T-DNA insertion events in transformation experiments with Silwet treatment (AS1–AS4)**.

The proportions of designated T-DNA insertion events in the respective transformation experiments are shown using the same structure as in Fig. 2C.

**Supplementary Figure 8. Proportions of designated T-DNA insertion events in transformation experiments using the B104 inbred line (B1–B8)**.

The proportions of designated T-DNA insertion events in the respective transformation experiments are shown using the same structure as in Fig. 2C.

**Supplementary Figure 9. Immature embryo and callus morphology of the B104 inbred line**.

A. Illustration comparing the embryonic callus-originating region between B104 and A188, based on top and side views of immature embryos. Dashed circles indicate the embryonic callus-originating region.

B. Size distribution of B104 embryos used in this study.

C. Examples of CoReCa morphology after the Sel-3 phase in B104 transformation experiments. Arrowheads indicate differently appearing CoReCa sectors originating from one callus. Bars = 1 mm.

**Supplementary Figure 10. GUS histochemical staining of regenerating plantlets from transformation experiments using hptII (hygromycin) and nptII (G418) selection systems**.

Small regenerating T0 plantlets from post-Reg1-2 calli from experiments AH1 and AH2 were examined by histochemical GUS staining. Arrowheads indicate positive GUS staining signals in regenerating roots.

**Supplementary Figure 11. Schematic representation of the crosses and screenings used for the isolation of mac1-2 and mac1-3 mutants**.

n.d., not detected.

**Supplementary Figure 12. Characterization of the mac1-2 and mac1-3 mutant phenotypes**.

A. Structure of the MAC1 locus showing guide RNA (gRNA) target sites and the isolated mac1-2 and mac1-3 mutant alleles.

B. Plant morphology of mac1-2 and mac1-3 compared with their respective wild-type (WT) siblings at the flowering stage. Bars = 1 m.

C. Anther morphology at the flowering stage. Bars = 1 mm.

D. Female fertility tested by crosses with wild-type pollen. Bars = 5 cm.

E. Bright-field images from confocal microscopy of early prophase I anthers, focusing on anther wall structure. Bars = 50 µm.

F. Schematic illustrations of early prophase I anthers in wild-type, mac1-2, and mac1-3 mutants, focusing on inner anther wall layer cells. Ep., epidermis; En., endothecium; Mi., middle layer; Ta., tapetum.

**Supplementary Figure 13. Schematic representation of the crosses and screenings used for the isolation of the acoz1-3 mutant, ACOZ1:YFP reporter, and establishment of the acoz1-3 complementation line carrying ACOZ1:YFP**.

A. Structure of the ACOZ1 locus showing guide RNA (gRNA) target sites, PCR primers used for screening (top), and examples of electropherograms from deletion mutant screening (bottom). Arrowheads indicate putative deletion-positive samples. M, DNA ladder marker (GeneRuler DNA Ladder Mix, Thermo Scientific).

B. The lineages used for isolation of the acoz1-3 mutant and ACOZ1:YFP reporter are shown on the left and right, respectively.

**Supplementary Figure 14. Characterization of the acoz1-3 mutant phenotype**.

A. Plant morphology of wild-type and acoz1-3 mutant plants at the flowering stage. Bars = 1 m.

B. Anther morphology at the flowering stage. Bars = 1 mm.

C. Pollen fertility assessed by Peterson’s staining. Bars = 100 µm.

D. Quantification of pollen fertility. Each bar represents the mean of individual plants determined from three independent anthers, and error bars indicate standard deviation.

E. Female fertility tested by crosses with wild-type pollen. Bars = 5 cm.

**Supplementary Figure 15. Characterization of SUN1:RFP localization in acoz1-3 mutant meiocytes**.

A. Expression pattern of SUN1:RFP in meiotic anthers compared with wild-type and acoz1-3 siblings. Bars = 50 µm.

B. Schematic representation of the image acquisition settings used for SUN1:RFP localization analysis.

C. Classification of SUN1:RFP localization patterns observed in zygotene(-like) meiocytes. Numbers correspond to the insets shown in Supplementary Fig. 15A.

D. Pie charts representing the proportions of SUN1:RFP localization patterns in zygotene(-like) meiocytes in acoz1-3 heterozygous (+/acoz1-3 het.; control) and acoz1-3 mutant plants.

**Supplementary Figure 16. Schematic representation of the transformation experiments, crosses, and screenings used to establish NCH:RFP reporters in the acoz1-3 mutant background**.

+/acoz1-3 het., plants heterozygous for acoz1-3.

**Supplementary Figure 17. Examples of NCH:RFP reporter expression in the acoz1-3 mutant background**.

Two additional NCH:RFP reporter events are shown.

A. Expression pattern of NCH2:RFP (ev.I, independent event I) in meiotic anthers compared with wild-type and acoz1-3 siblings. Bars = 50 µm.

B. Expression pattern of NCH1:RFP (ev.D) in meiotic anthers compared with wild-type and acoz1-3 siblings. Bars = 50 µm.

## References

1. Aesaert, S., Impens, L., Coussens, G. et al. (2022) Optimized Transformation and Gene Editing of the B104 Public Maize Inbred by Improved Tissue Culture and Use of Morphogenic Regulators. Front Plant Sci 13, 883847.

2. Armstrong, C.L., Green, C.E. and Phillips, R.L. (1991) Development and availability of germplasm with high Type II culture formation response.

3. Brettschneider, R., Becker, D. and Lörz, H. (1997) Efficient transformation of scutellar tissue of immature maize embryos. Theoretical and Applied Genetics 94, 737–748.

4. Char, S.N., Neelakandan, A.K., Nahampun, H. et al. (2017) An Agrobacterium-delivered CRISPR/Cas9 system for high-frequency targeted mutagenesis in maize. Plant Biotechnol J 15, 257–268.

5. Clough, S.J. and Bent, A.F. (1998) Floral dip: a simplified method for Agrobacterium-mediated transformation of Arabidopsis thaliana. Plant J 16, 735–743.

6. Curtis, I.S. and Nam, H.G. (2001) Transgenic radish (Raphanus sativus L. longipinnatus Bailey) by floral-dip method–plant development and surfactant are important in optimizing transformation efficiency. Transgenic research 10, 363–371.

7. Dellaporta, S.L., Wood, J. and Hicks, J.B. (1983) A plant DNA minipreparation: version II. Plant molecular biology reporter

8. Fauser, F., Schiml, S. and Puchta, H. (2014) Both CRISPR/Cas-based nucleases and nickases can be used efficiently for genome engineering in Arabidopsis thaliana. Plant J 79, 348–359.

9. Feng, C., Su, H., Bai, H. et al. (2018) High-efficiency genome editing using a dmc1 promoter-controlled CRISPR/Cas9 system in maize. Plant Biotechnol J 16, 1848–1857.

10. Ge, F., Qu, J., Liu, P., Pan, L., Zou, C., Yuan, G., Yang, C., Pan, G., Huang, J. and Ma, L. (2022) Genome assembly of the maize inbred line A188 provides a new reference genome for functional genomics. The Crop Journal 10, 47–55.

11. Gelvin, S.B. (2010) Plant proteins involved in Agrobacterium-mediated genetic transformation. Annual review of phytopathology 48, 45–68.

12. Han, L. and Cai, H. (2024) Progress on Genetic Transformation of Sorghum. Scientia Agricultura Sinica 57, 454–468.

13. He, Y., Zhang, T., Sun, H., Zhan, H. and Zhao, Y. (2020) A reporter for noninvasively monitoring gene expression and plant transformation. Hortic Res 7, 152.

14. Hernandes-Lopes, J., Pinto, M.S., Vieira, L.R. et al. (2023) Enabling genome editing in tropical maize lines through an improved, morphogenic regulator-assisted transformation protocol. Front Genome Ed 5, 1241035.

15. Hwang, H.-H., Yu, M. and Lai, E.-M. (2017) Agrobacterium-mediated plant transformation: biology and applications. The arabidopsis book 15, e0186.

16. Ishida, Y., Hiei, Y. and Komari, T. (2007) Agrobacterium-mediated transformation of maize. Nat Protoc 2, 1614–1621.

17. Iwase, A., Kondo, Y., Laohavisit, A. et al. (2021) WIND transcription factors orchestrate wound-induced callus formation, vascular reconnection and defense response in Arabidopsis. New Phytol 232, 734–752.

18. Jing, J., Wu, N., Xu, W., Wang, Y. and Pawlowski, W.P. (2022) An F-box protein ACOZ1 functions in crossover formation by ensuring proper chromosome compaction during maize meiosis. New …

19. Kausch, A.P., Wang, K., Kaeppler, H.F. and Gordon-Kamm, W. (2021) Maize transformation: history, progress, and perspectives. Molecular Breeding

20. Komari, T., Halperin, W. and Nester, E.W. (1986) Physical and functional map of supervirulent Agrobacterium tumefaciens tumor-inducing plasmid pTiBo542. J Bacteriol 166, 88–94.

21. Li, J., Ye, X., An, B., Du, L. and Xu, H. (2012) Genetic transformation of wheat: current status and future prospects. Plant Biotechnology Reports 6, 183–193.

22. Lin, G., He, C., Zheng, J., Koo, D.H., Le, H. and Zheng, H. (2021) Chromosome-level genome assembly of a regenerable maize inbred line A188. Genome biology

23. Liu, S.-J., Wei, Z.-M. and Huang, J.-Q. (2008) The effect of co-cultivation and selection parameters on Agrobacterium-mediated transformation of Chinese soybean varieties. Plant cell reports 27, 489–498.

24. Lowe, K., La Rota, M., Hoerster, G., Hastings, C., Wang, N., Chamberlin, M., Wu, E., Jones, T. and Gordon-Kamm, W. (2018) Rapid genotype “independent” Zea mays L. (maize) transformation via direct somatic embryogenesis. In Vitro Cell Dev Biol Plant 54, 240–252.

25. Lowe, K., Wu, E., Wang, N. et al. (2016) Morphogenic Regulators Baby boom and Wuschel Improve Monocot Transformation. Plant Cell 28, 1998–2015.

26. Marand, A.P., Jiang, L., Gomez-Cano, F. and Minow, M.A.A. (2025) The genetic architecture of cell type–specific cis regulation in maize. Science

27. Miller, M., Tagliani, L., Wang, N., Berka, B. and Bidney, D. (2002) High Efficiency Transgene Segregation in Co-Transformed Maize Plants using an Agrobacterium Tumefaciens 2 T-DNA Binary System. Transgenic *…*

28. Mookkan, M., Nelson-Vasilchik, K., Hague, J., Kausch, A. and Zhang, Z.J. (2018) Morphogenic Regulator-Mediated Transformation of Maize Inbred B73. Curr Protoc Plant Biol 3, e20075.

29. Peterson, R., Slovin, J.P. and Chen, C. (2010) A simplified method for differential staining of aborted and non-aborted pollen grains. International Journal of Plant …

30. Rahnama, H. (2010) Agrobacterium mediated transformation of maize (Zea mays L.). Journal of Sciences

31. Raji, J.A., Frame, B., Little, D., Santoso, T.J. and Wang, K. (2018) Agrobacterium– and Biolistic-Mediated Transformation of Maize B104 Inbred. Methods Mol Biol 1676, 15–40.

32. Rojano-Delgado, A.M., Priego-Capote, F., De Prado, R. and De Castro, M.D.L. (2014) Qualitative/quantitative strategy for the determination of glufosinate and metabolites in plants. Analytical and bioanalytical chemistry 406, 611.

33. Sheridan, W.F., Avalkina, N.A., Shamrov, I.I., Batygina, T.B. and Golubovskaya, I.N. (1996) The mac1 gene: controlling the commitment to the meiotic pathway in maize. Genetics 142, 1009–1020.

34. Singh, R.K. and Prasad, M. (2016) Advances in Agrobacterium tumefaciens-mediated genetic transformation of graminaceous crops. Protoplasma 253, 691.

35. Sjahril, R. and Mii, M. (2006) High-efficiency Agrobacterium-mediated transformation of Phalaenopsis using meropenem, a novel antibiotic to eliminate Agrobacterium. The Journal of Horticultural Science and Biotechnology 81, 458–464.

36. Springer, W.D., Green, C.E. and Kohn, K.A. (1979) A histological examination of tissue culture initiation from immature embryos of maize. Protoplasma

37. Staut, J., Pérez, N.M., Ferrando, A.M. and Dissanayake, I. (2025) A map of integrated cis-regulatory elements enhances gene-regulatory analysis in maize. Plant *…*

38. Vasil, V., Lu, C.Y. and Vasil, I.K. (1985) Histology of somatic embryogenesis in cultured immature embryos of maize (Zea mays L.). Protoplasma

39. Vega, J.M., Yu, W., Kennon, A.R., Chen, X. and Zhang, Z.J. (2008) Improvement of Agrobacterium-mediated transformation in Hi-II maize (Zea mays) using standard binary vectors. Plant Cell Rep 27, 297–305.

40. Vos, P., Hogers, R., Bleeker, M., Reijans, M., Lee, T.V.D., Hornes, M., Friters, A., Pot, J., Paleman, J. and Kuiper, M. (1995) AFLP: a new technique for DNA fingerprinting. Nucleic acids research 23, 4407–4414.

41. Wang, C.J., Nan, G.L., Kelliher, T., Timofejeva, L., Vernoud, V., Golubovskaya, I.N., Harper, L., Egger, R., Walbot, V. and Cande, W.Z. (2012) Maize multiple archesporial cells 1 (mac1), an ortholog of rice TDL1A, modulates cell proliferation and identity in early anther development. Development 139, 2594–2603.

42. Wang, N., Arling, M., Hoerster, G., Ryan, L. and Wu, E. (2020) An efficient gene excision system in maize. Frontiers in plant …

43. Xi, J., Patel, M., Dong, S., Que, Q. and Qu, R. (2018) Acetosyringone treatment duration affects large T-DNA molecule transfer to rice callus. BMC biotechnology

44. Yuan, X., Cai, B., Hamamura, Y., Schnittger, A. and Yang, C. (2025) SCF^RMF^-dependent degradation of the nuclear lamina releases the somatic chromatin mobility restriction for meiotic recombination. Science Advances

45. Zhang, F., Tang, D., Shen, Y., Xue, Z., Shi, W. and Ren, L. (2017) The F-box protein ZYGO1 mediates bouquet formation to promote homologous pairing, synapsis, and recombination in rice meiosis. The Plant …

46. Zhao, Z.Y., Gu, T., Cai, T., Tagliani, L.A., Hondred, D., Bond, D., Krell, S., Rudert, M.L., Bruce, W.B. and Pierce, D.A. (1998) Molecular analysis of T0 plants transformed by Agrobacterium and comparison of Agrobacterium-mediated transformation with bombardment transformation in maize.

