## Supplemental table and figures for "Reconstructing tissue culture to improve Agrobacterium-mediated transformation of maize"

**Supplementary Table 1: Summary of constructs generated and used in this study.**

| Experiment ID | Type | Represetative costruct name | Binary vector backbone (selection marker gene) | Target locus (loci) for CRISPR/Cas9, or Genes cloned for Reporter | Gene symbols, (appeared on MaizeGDB B73v5 annotation) | gRNA sequece 1 | gRNA sequece 2 | gRNA sequece 3 | gRNA sequece 4 | Cloned Genomic region for reporter construct (in the B73v5 corrdinates) |
|---|---|---|---|---|---|---|---|---|---|---|
| A1 | Reporter | UBIpro:RUBY_DMC1pro:SpCas9_no_gRNA/p7oM | p7oM-LH (bar) | - | - | - | - | - | - | - |
| A2 | Reporter | PAIR1:YFP/p7oM | p7oM-LH (bar) | Zm00001eb404350 | prd3 | - | - | - | - | chr9:161344704-161336350 |
| A3 | Reporter | PAIR1:RFP/p7oM | p7oM-LH (bar) | Zm00001eb404350 | prd3 | - | - | - | - | chr9:161344704-161336350 |
| A4 | Reporter | ASY1:YFP/p7oM | p7oM-LH (bar) | Zm00001eb102420 | asy1 | - | - | - | - | chr2:198050067-198065001 (2 TE sequences removed) |
| A5 | Reporter | ASY1:RFP/p7oM | p7oM-LH (bar) | Zm00001eb102420 | asy1 | - | - | - | - | chr2:198050067-198065001 (2 TE sequences removed) |
| A6 | CRISPR/Cas9 | UBIpro:SaCas9_gRNA13/p7oM | p7oM-LH (bar) | Zm00001eb045380 | dcl102 | TGAAGAGATTTATTGGCCAGATGAAT | CCATGTGACAGAGGCCACAGATGGAT | - | - | - |
| A7 | Reporter | YFP:PAIR1/p7oM | p7oM-LH (bar) | Zm00001eb404350 | prd3 | - | - | - | - | chr9:161344704-161336350 |
| A8 and A9 | CRISPR/Cas9 | UBIpro:SpCas9_gRNA23/p7oM | p7oM-LH (bar) | Zm00001eb145140 | zip4 | CGAGGAGGAGGAGGAAGCAGCGG | TGCACTACTGAACTTATGTACGG | - | - | - |
| A10 | CRISPR/Cas9 | UBIpro:SaCas9_gRNA10/p7oM | p7oM-LH (bar) | Zm00001eb132020 | cl38706_1 | GTACTGTGCTATTAATTATCAAAGAAT | GAGTGGGGCAGATAAACACAAATGAGT | - | - | - |
| A11 | CRISPR/Cas9 | UBIpro:SaCas9_gRNA32B/p7oM | p7oM-LH (bar) | Zm00001eb408950 | mac1 | GGCTTCGGCGCCTGCATCACAACCGGGT | GTGCGGCCGAGCCGGATGGACGAGGGGT | - | - | - |
| ASt-a and ASt-b | Reporter | UBIpro:GUS/p7oM | p7oM-LH (bar) | - | | - | - | - | - | - |
| AS1 | CRISPR/Cas9 | UBIpro:SaCas9_gRNA29A/p7oM | p7oM-LH (bar) | Zm00001eb233800 | dsy2 | GTCTTTTTATTATTTACATCGACAGAAT | CTGAATTGCAGGAGTTCTCTTGAGGAT | - | - | - |
| AS2 | CRISPR/Cas9 | UBIpro:SaCas9_gRNA29B/p7oM | p7oM-LH (bar) | Zm00001eb233800 | dsy2 | GACTATATGTCAGCACCTTGAGGGGGT | GCTAAATGTGGTTCTTCAAATGCAGGAT | - | - | - |
| AS3 | Reporter | UBIpro:RUBY/p7oM | p7oM-LH (bar) | - | - | - | - | - | - | - |
| AS4 | CRISPR/Cas9 | UBIpro:SaCas9_gRNA28/p7oM | p7oM-LH (bar) | Zm00001eb423930 | zyp1 | ATTAGCTTCAGAACTACAAGGGAGAAT | ATGTTGAACCATAACAATTTCTTGAGT | GATCAACCAGCTTCGTAGTGTTCTGGAT | TCTTGCTGCATATACAATACCATGAGT | - |
| B1 and B7 | CRISPR/Cas9 | UBIpro:SaCas9_gRNA6/p7oM | p7oM-LH (bar) | Zm00001eb274370, Zm00001eb195020, Zm00001eb025660 | IDP4036, lbd25 | TTCACTTGCAAGTGTCCCGTCAGGAGT | GAATGCATGTCTTCCTCTATGTTTGAGT | - | - | - |
| B2 and B8 | CRISPR/Cas9 | UBIpro:SaCas9_gRNA13/p7oM | p7oM-LH (bar) | Zm00001eb045380 | dcl102 | TGAAGAGATTTATTGGCCAGATGAAT | CCATGTGACAGAGGCCACAGATGGAT | - | - | - |
| B3 | CRISPR/Cas9 | UBIpro:SpCas9_gRNA17/p7oM | p7oM-LH (bar) | Zm00001eb160270 | rad51c | AATCAATGTACAAATCCCAGTGG | GTTACCCATCATGCGTATGAGGGG | - | - | - |
| B4 | CRISPR/Cas9 | UBIpro:RUBY_DMC1pro:SpCas9_gRNA17/p7oM | p7oM-LH (bar) | Zm00001eb160270 | rad51c | AATCAATGTACAAATCCCAGTGG | GTTACCCATCATGCGTATGAGGGG | - | - | - |
| B5 | CRISPR/Cas9 | UBIpro:SpCas9_gRNA18/p7oM | p7oM-LH (bar) | Zm00001eb362590 | mlh1 | GTTCCTTTAACTGCAGCACAGGG | GCTGTGACATAAGAAGCAAATGG | - | - | - |
| B6 | CRISPR/Cas9 | UBIpro:RUBY_DMC1pro:SpCas9_gRNA18/p7oM | p7oM-LH (bar) | Zm00001eb362590 | mlh1 | GTTCCTTTAACTGCAGCACAGGG | GCTGTGACATAAGAAGCAAATGG | - | - | - |
| AH1 | Reporter | UBIpro:GUS/p6oM | p6oM-LH (hptII) | - | - | - | - | - | - | - |
| AH2 | CRISPR/Cas9 | UBIpro:SaCas9_gRNA32B/p6oM | p6oM-LH (hptII) | Zm00001eb408950 | mac1 | GGCTTCGGCGCCTGCATCACAACCGGGT | GTGCGGCCGAGCCGGATGGACGAGGGGT | - | - | - |
| AG1 | Reporter | UBIpro:GUS/p9oM | p9oM-LH (nptII) | - | - | - | - | - | - | - |
| AG2 | CRISPR/Cas9 | UBIpro:SaCas9_gRNA32B/p9oM | p9oM-LH (nptII) | Zm00001eb408950 | mac1 | GGCTTCGGCGCCTGCATCACAACCGGGT | GTGCGGCCGAGCCGGATGGACGAGGGGT | - | - | - |
| | | | | | | | | - | - | |
| ACOZ1-CRISPR | CRISPR/Cas9 | UBIpro:SaCas9_gRNA7 | p7oM-LH (bar) | Zm00001eb119810 | acoz1 | CTGCATTGAGAAATACTGGCTCAGGAT | GAAGTAAGCGCAGCGAGATTGTAGGAT | - | - | - |
| ACOZ1-reporter | Reporter | ACOZ1:YFP/p7oM | p7oM-LH (bar) | Zm00001eb119810 | acoz1 | - | - | - | - | chr3:3165481-3173522 |
| SUN1-reporter | Reporter | SUN1:RFP/p7oM | p7oM-LH (bar) | Zm00001eb233650 | sun1 | - | - | - | - | chr5:90720207-90730914 |
| NCH1-reporter | Reporter | NCH1:RFP/p7oM | p7oM-LH (bar) | Zm00001eb188670 | nch1 | - | - | - | - | chr4:166234321-166243285 |
| NCH2-reporter | Reporter | NCH2:RFP/p7oM | p7oM-LH (bar) | Zm00001eb152080 | nch2 | - | - | - | - | chr3:199092297-199111748 (1 TE sequence removed) |

PAM sequences of gRNA were underlined.

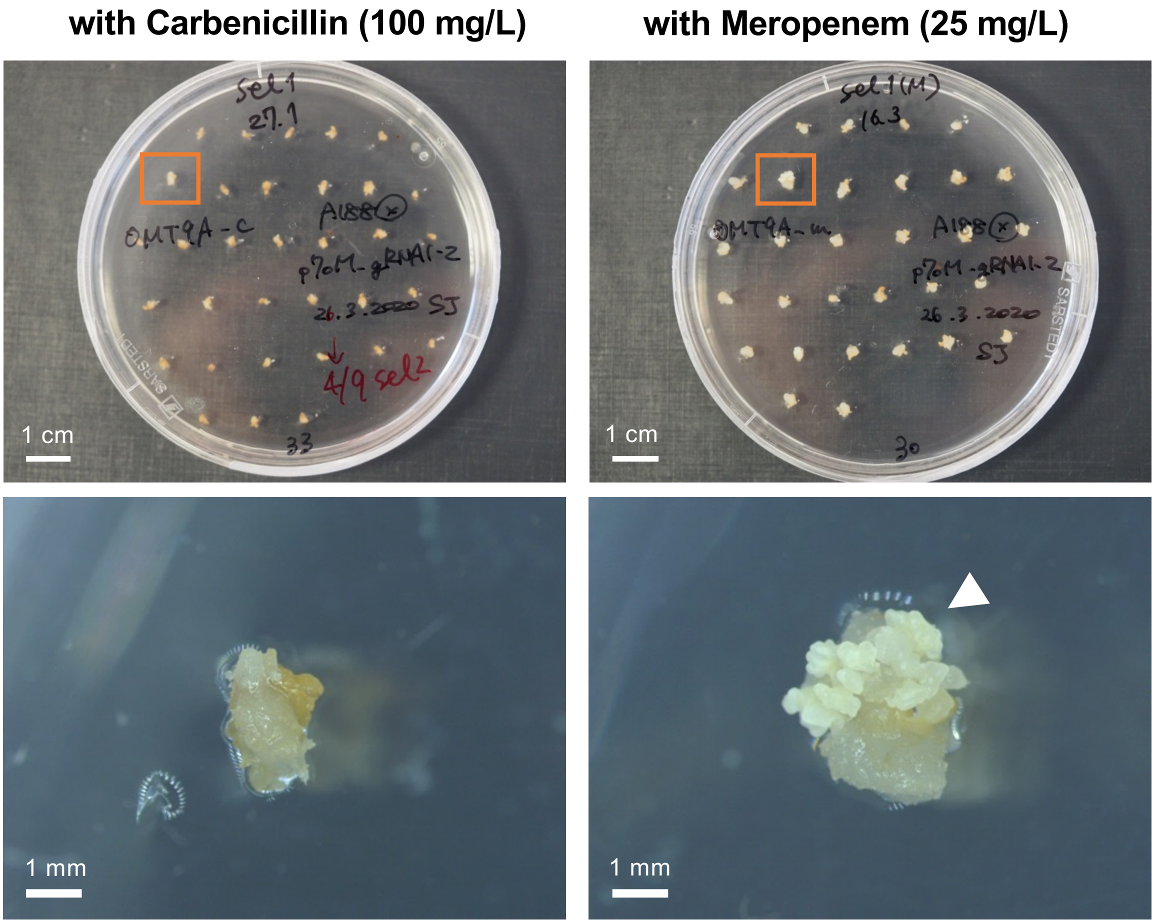


### Supplementary figure 1

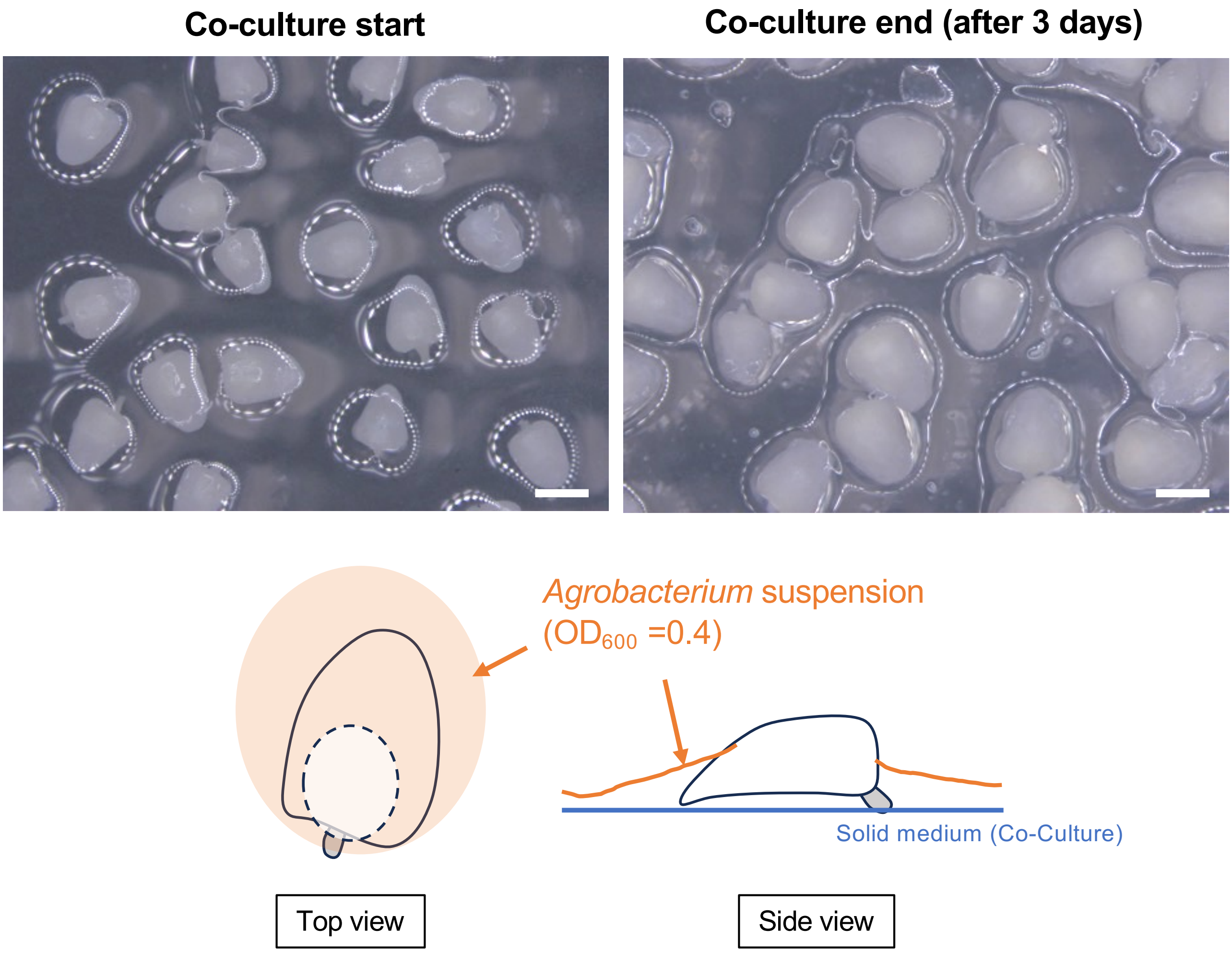


### Supplementary figure 2

**without embedding** **with embedding**

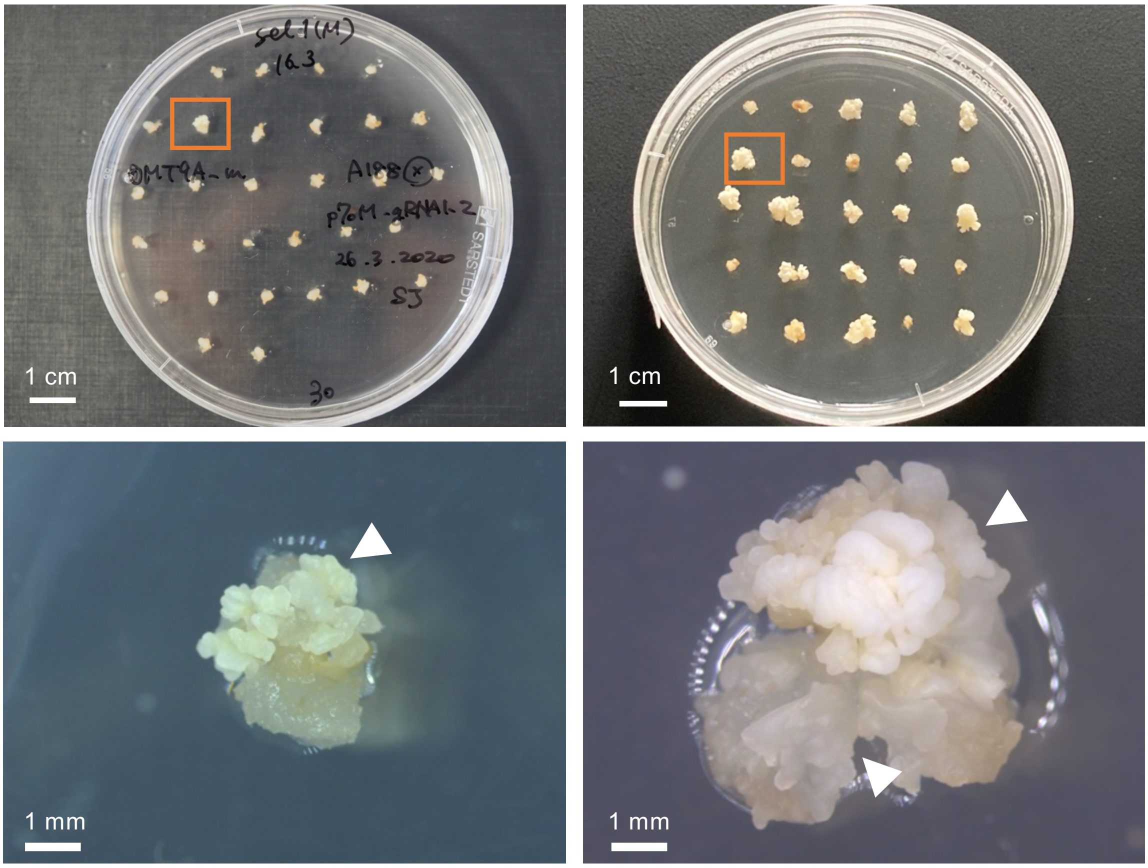


### Supplementary figure 3

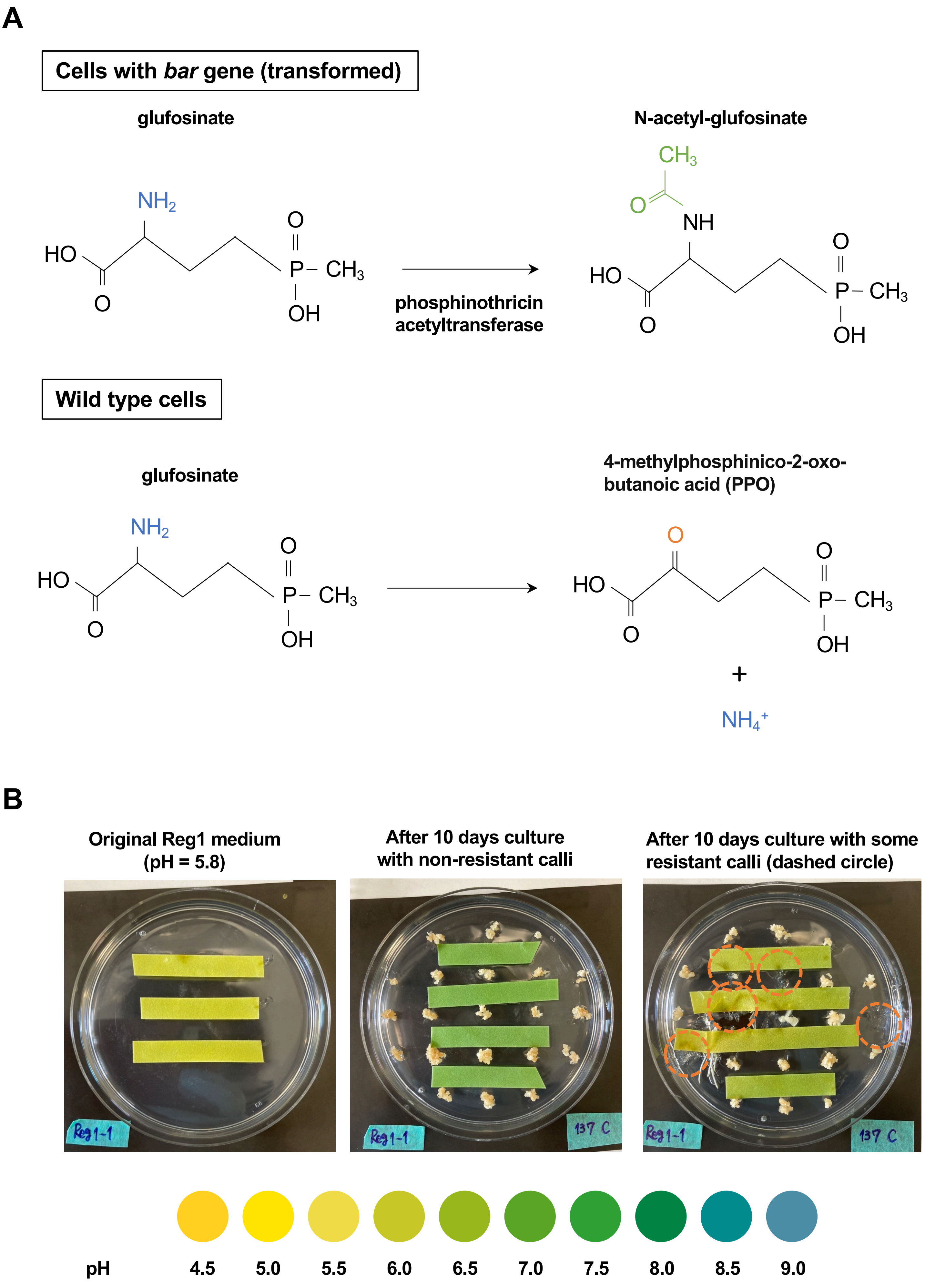


### Supplementary figure 4

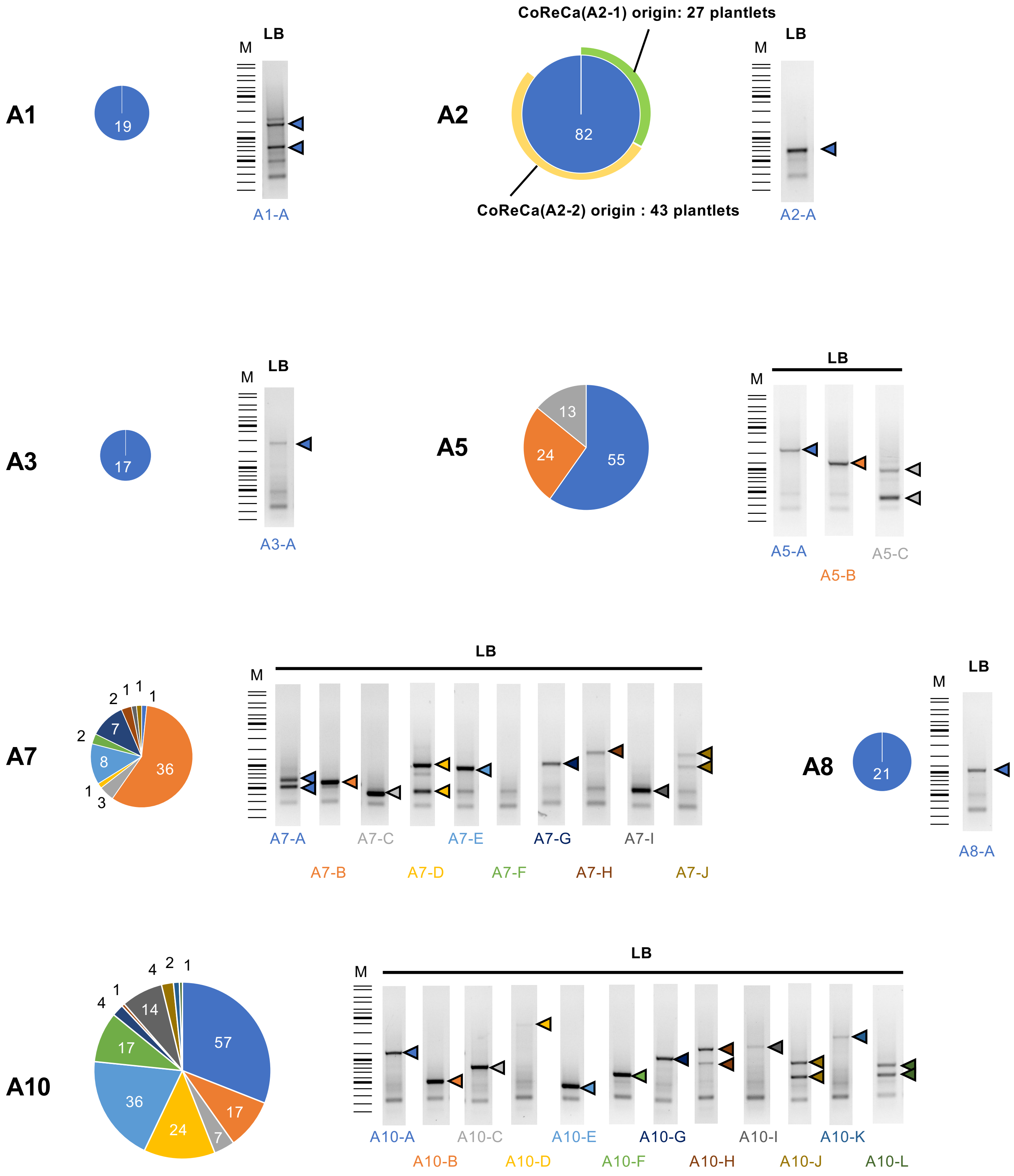


### Supplementary figure 5

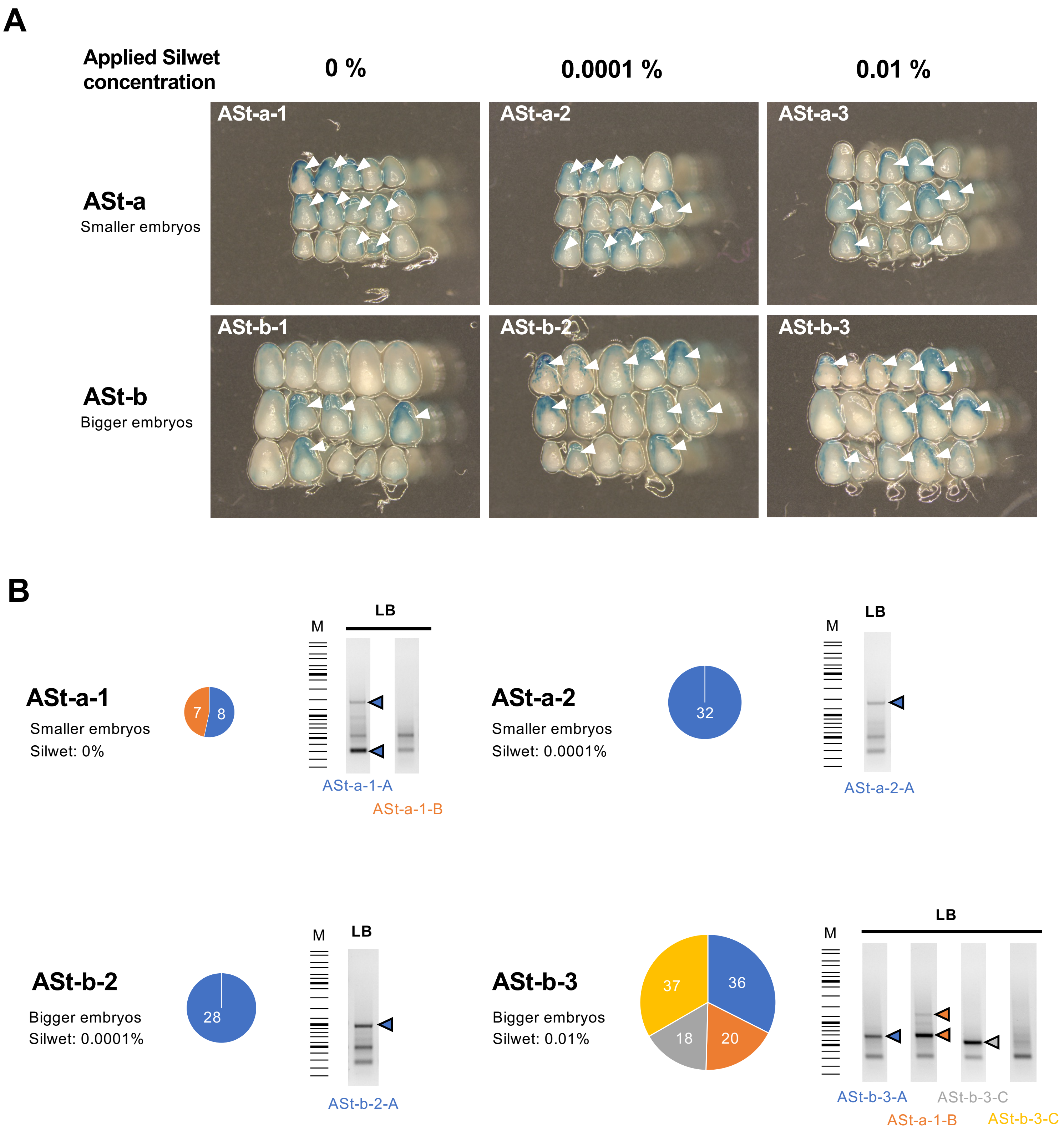


### Supplementary figure 6

AS1

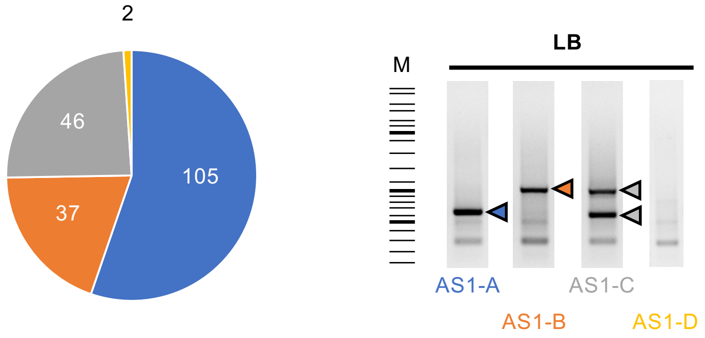


AS2

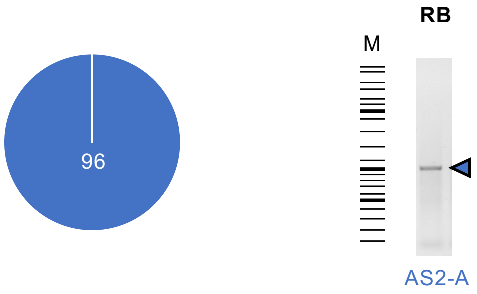


AS3

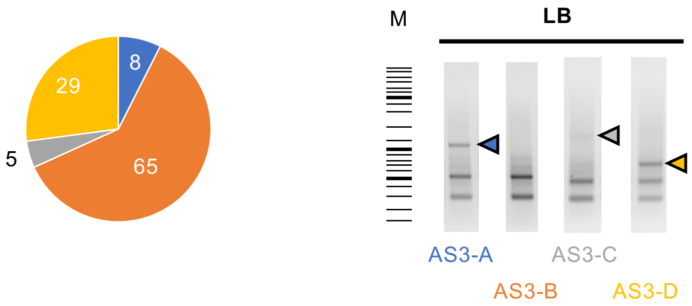


AS4

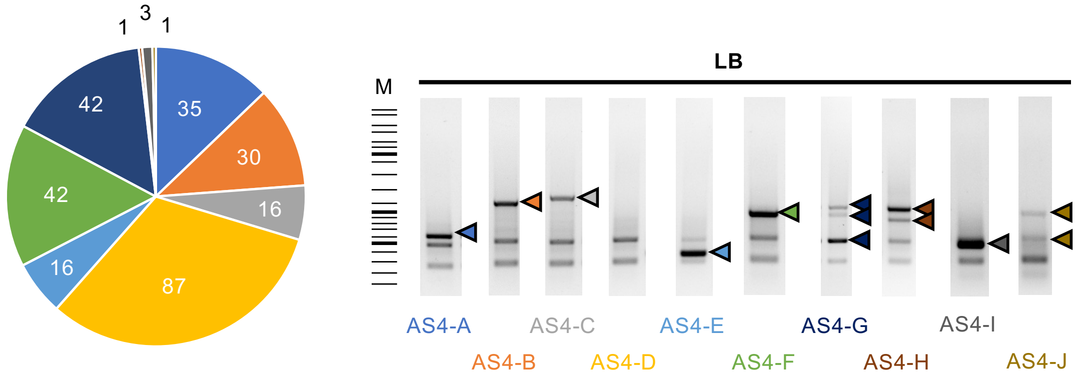


### Supplementary figure 7

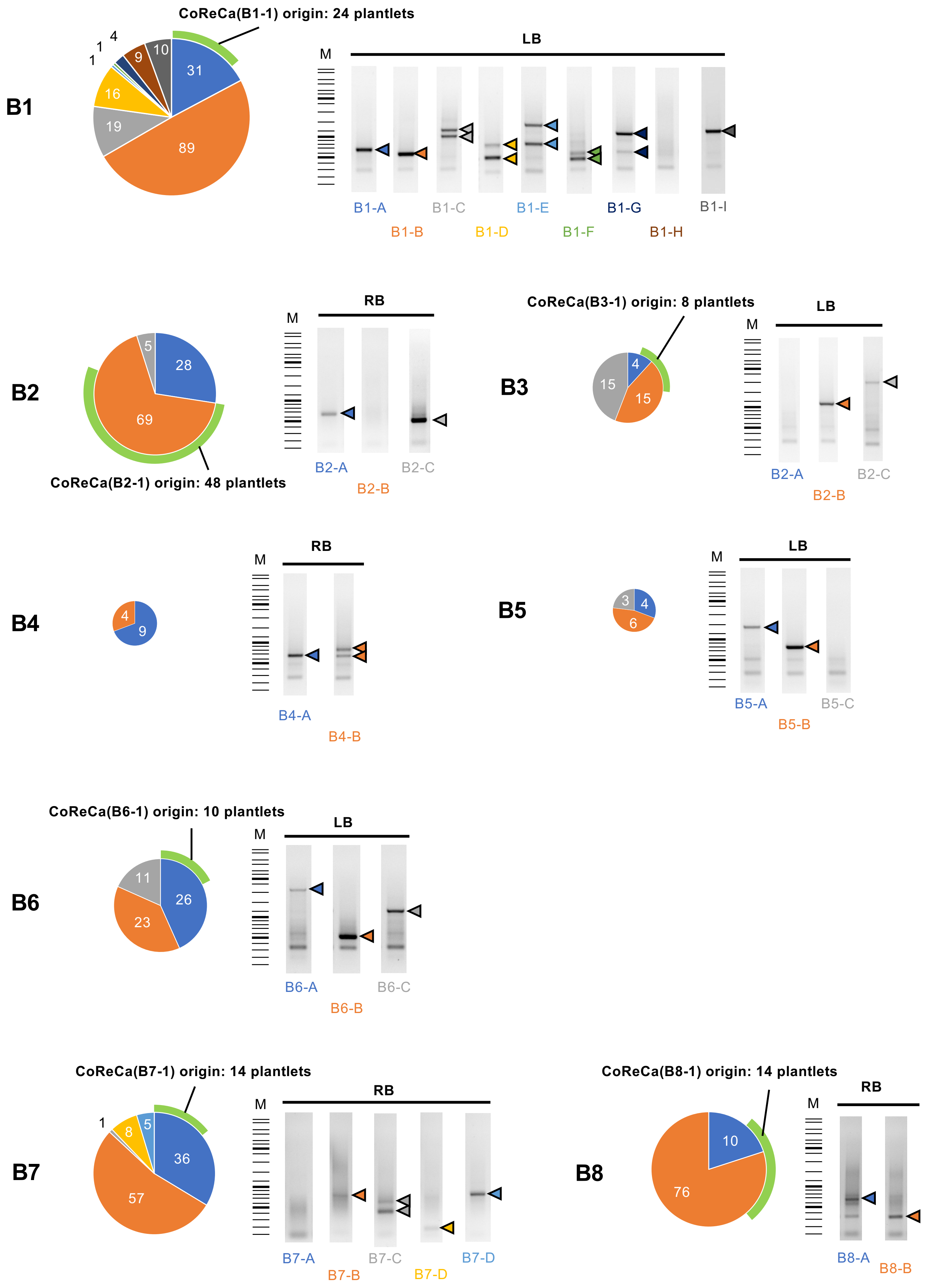


### Supplementary figure 8

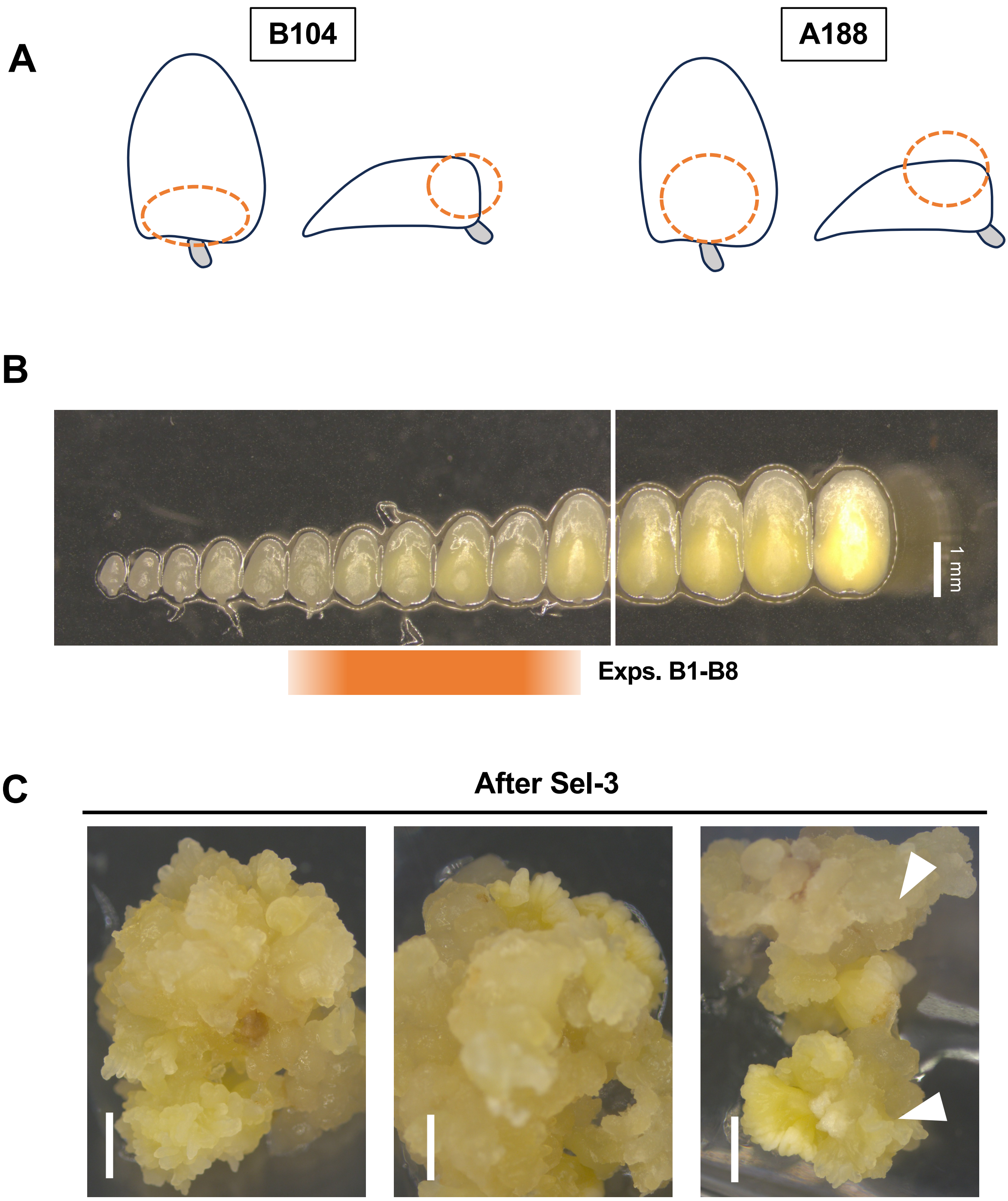


### Supplementary figure 9

AH1 (hygromycin) AG1 (G418)

T0 plantlets

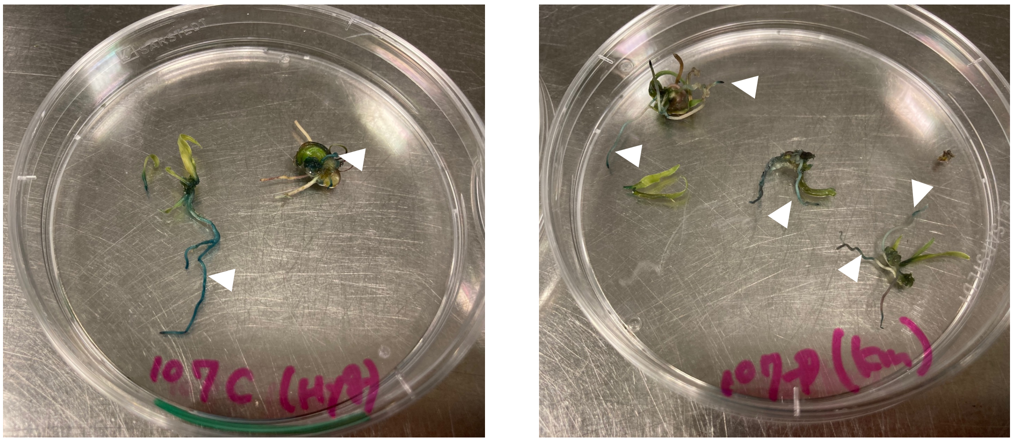

**Supplementary figure 10**

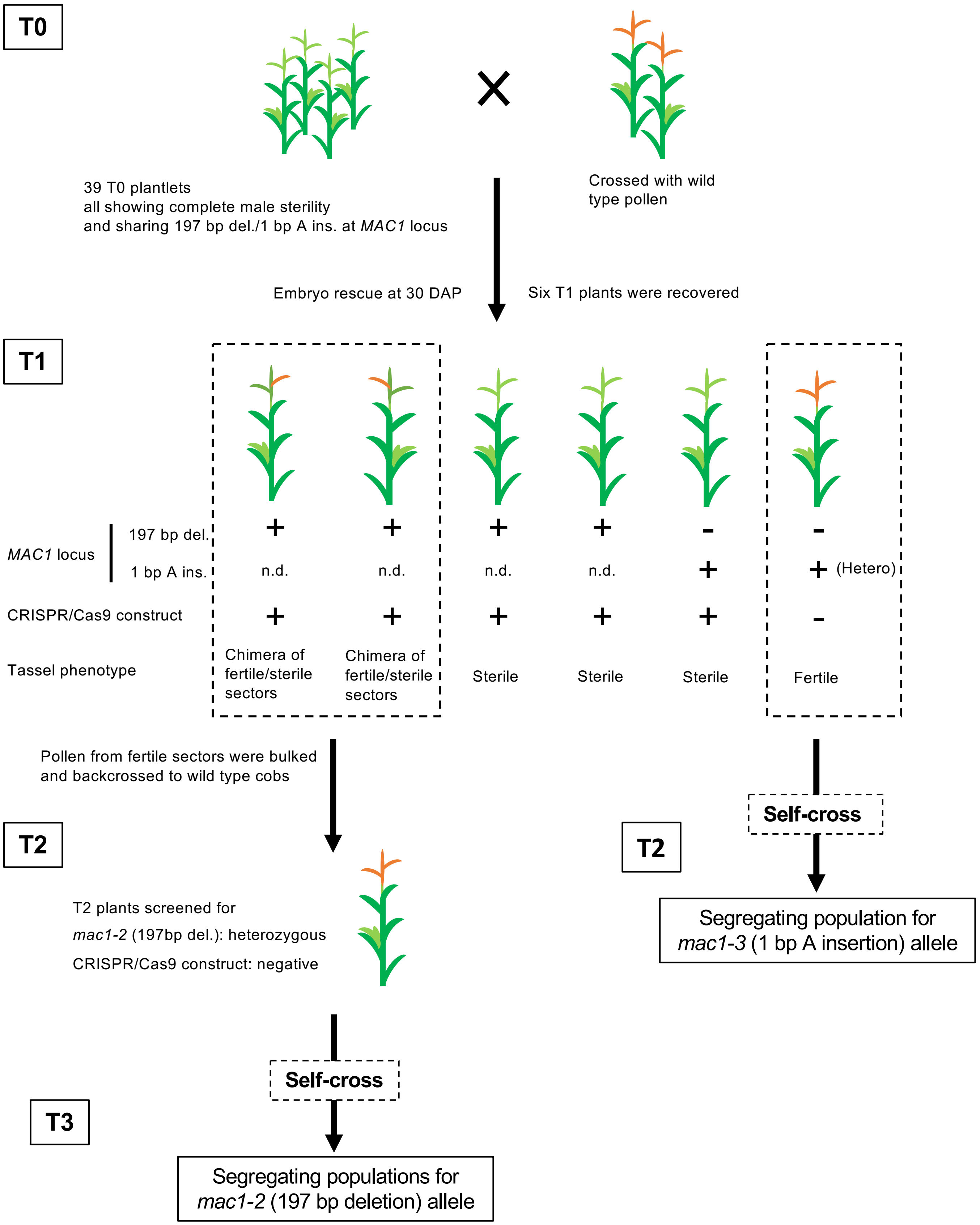


### Supplementary figure 11

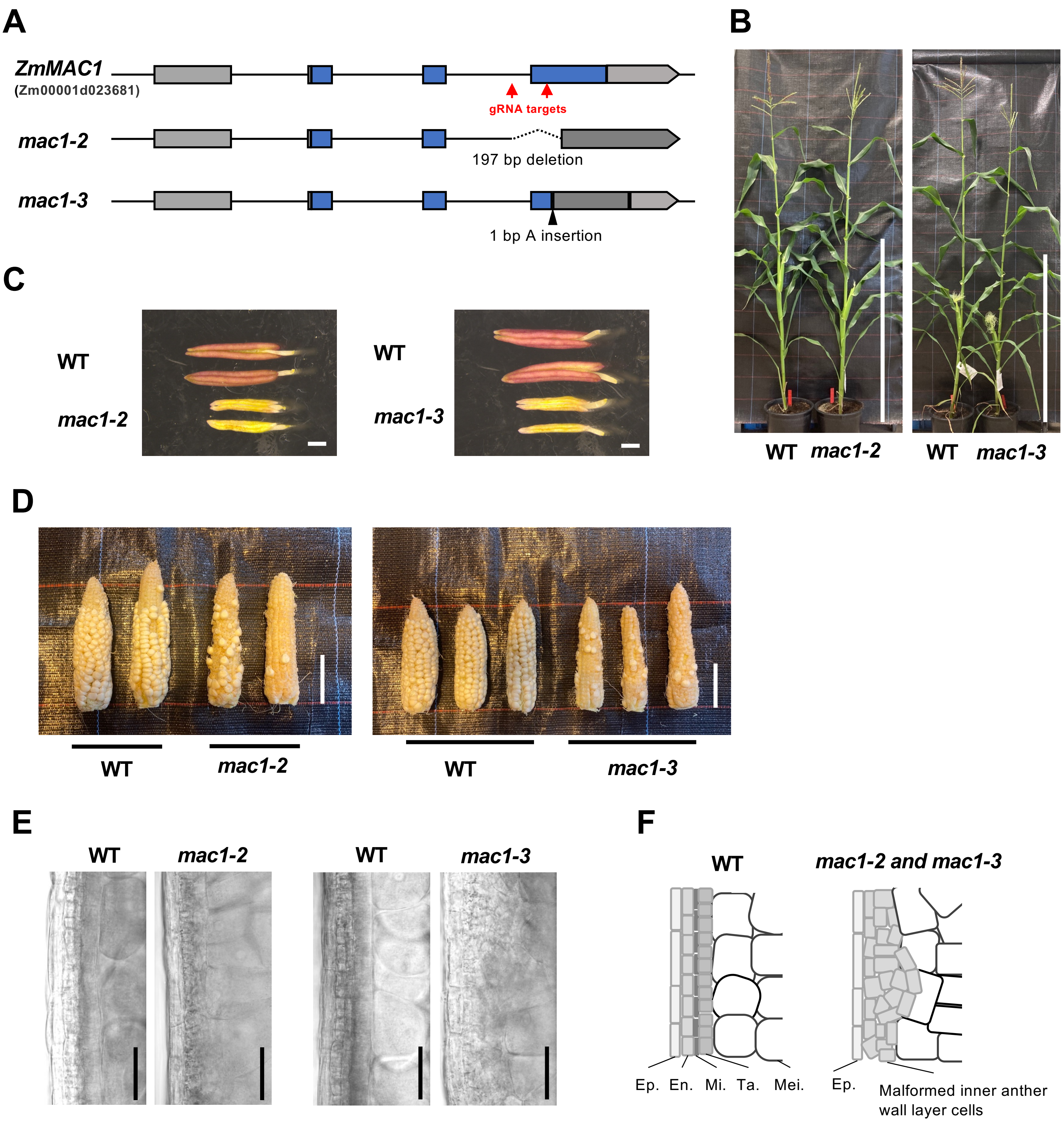


### Supplementary figure 12

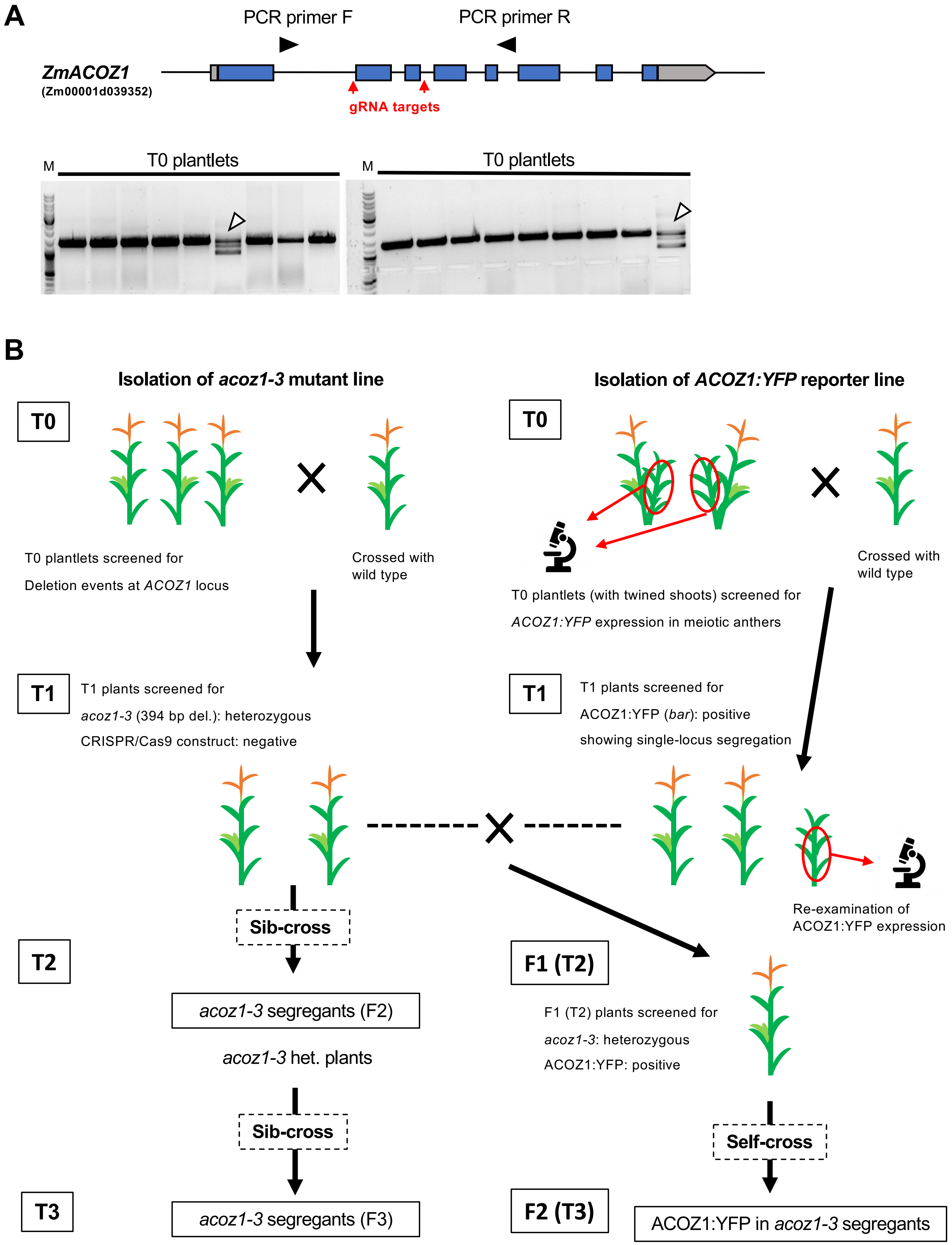


#### Supplementary figure 13

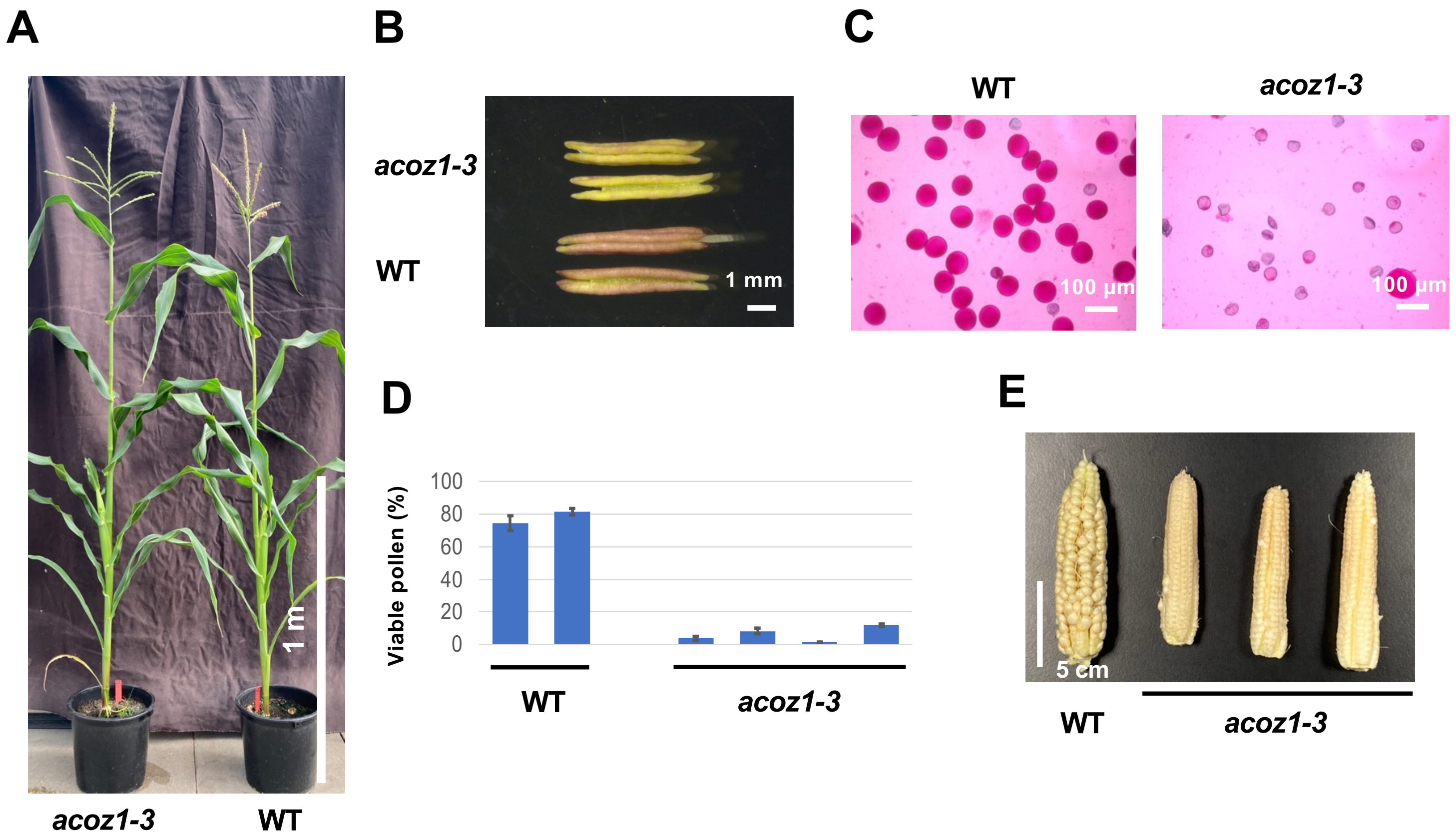


**Supplementary figure 14**

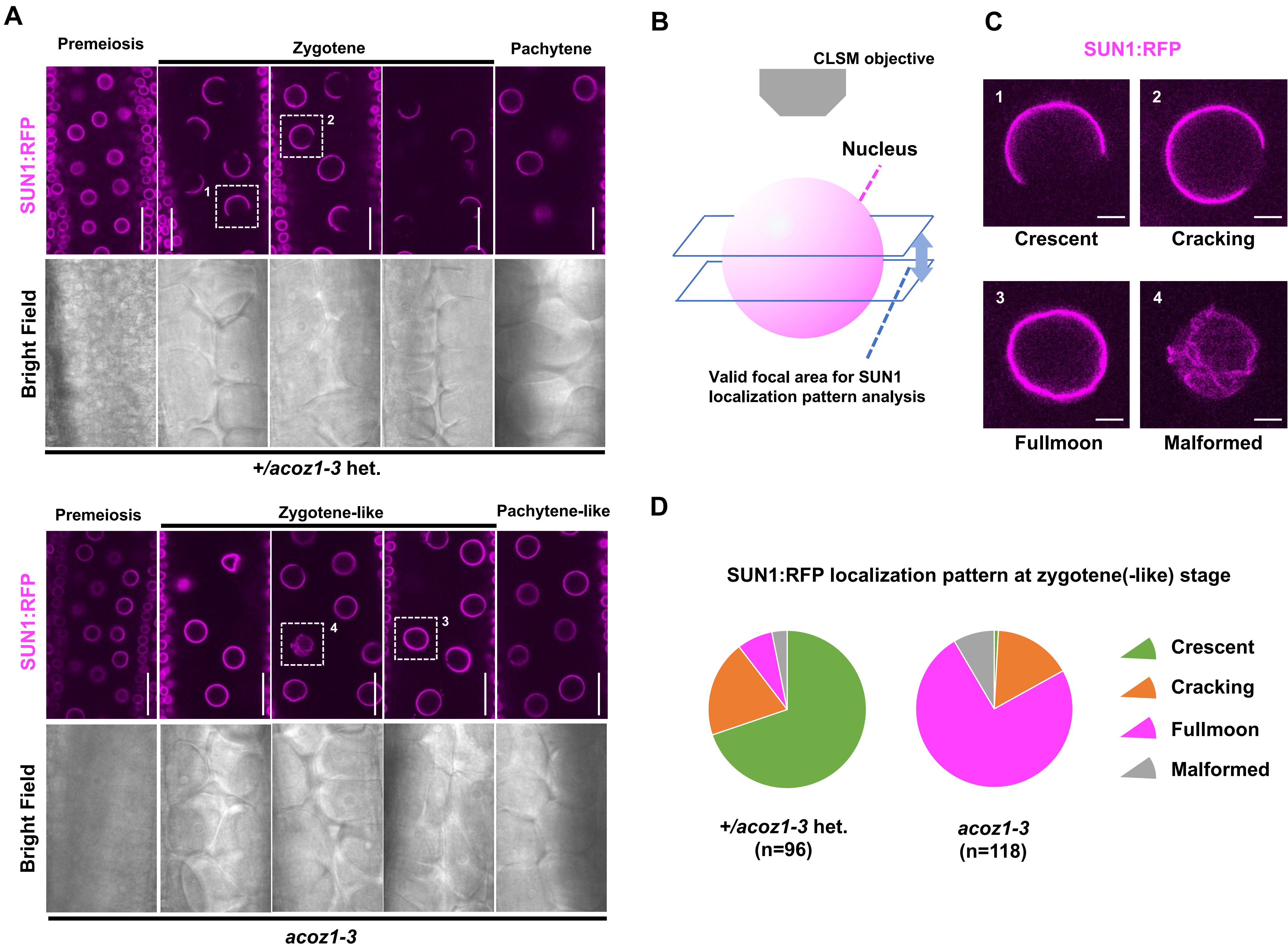


**Supplementary figure 15**

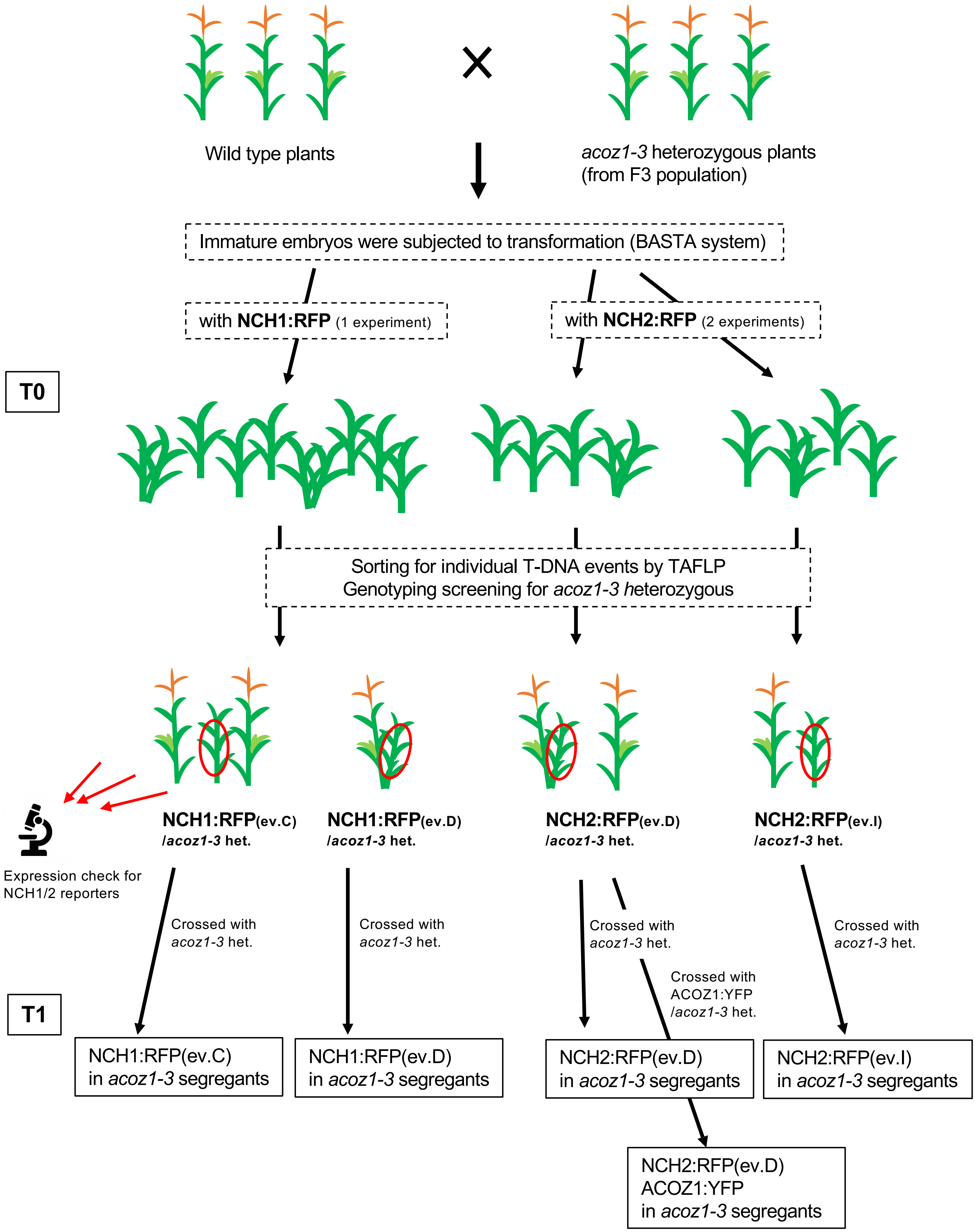


### Supplementary figure 16

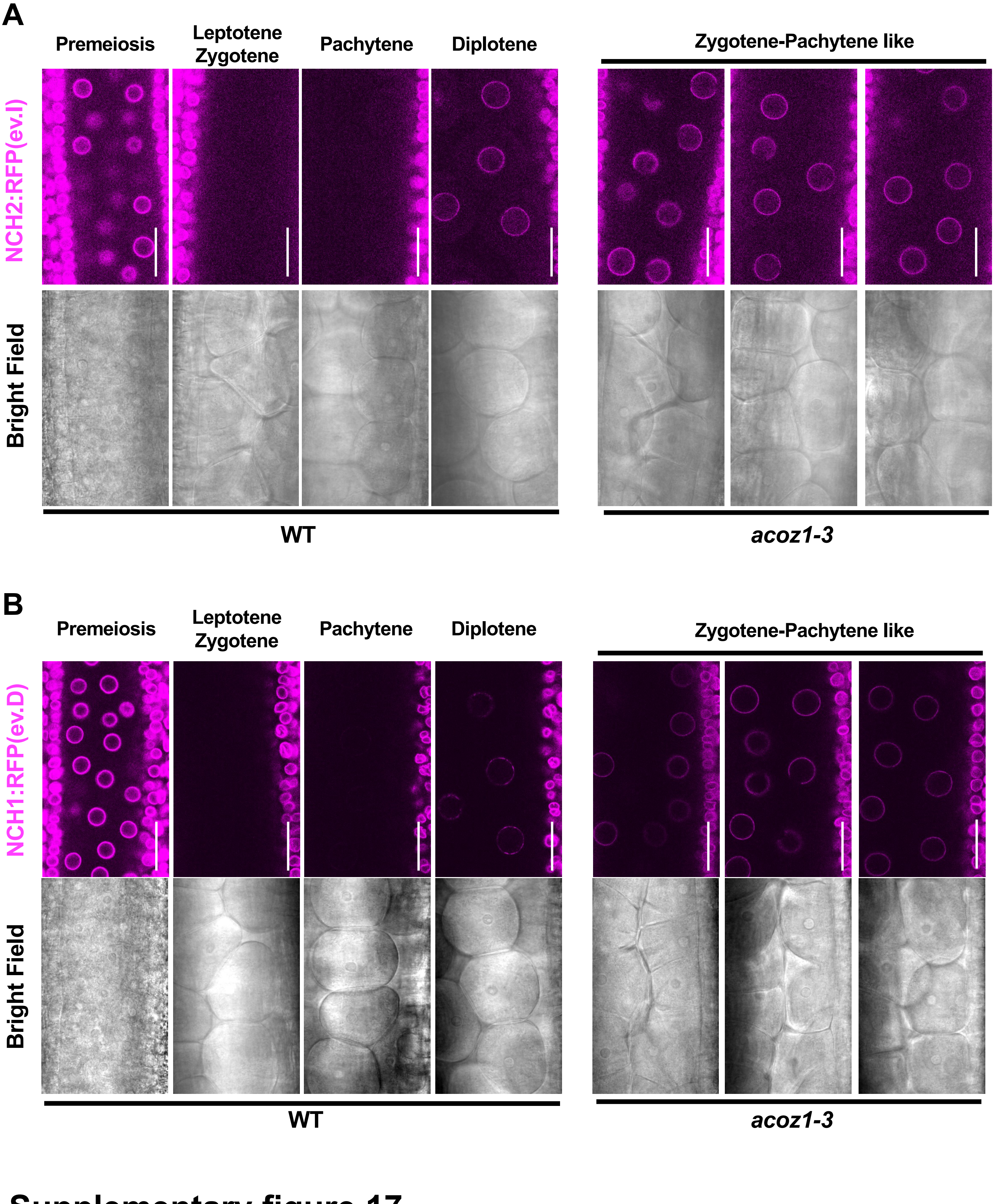


**Supplementary figure 17**
